# Mapping the semi-nested community structure of 3D chromosome contact networks

**DOI:** 10.1101/2022.06.24.497560

**Authors:** Dolores Bernenko, Sang Hoon Lee, Per Stenberg, Ludvig Lizana

**Affiliations:** Department of Physics, Integrated Science Lab, Umeå University, SE-901 87 Umeå, Sweden; Department of Physics and Research Institute of Natural Science, Gyeongsang National University, Jinju 52828, Korea and Future Convergence Technology Research Institute, Gyeongsang National University, Jinju 52849, Korea; EMG, Umeå University, SE-901 87 Umeå, Sweden; Integrated Science Lab, Department of Physics, Umeå University, SE-901 87 Umeå, Sweden

## Abstract

Mammalian DNA folds into 3D structures that facilitate and regulate genetic processes such as transcription, DNA repair, and epigenetics. Several insights derive from chromosome capture methods, such as Hi-C, which allow researchers to construct contact maps depicting 3D interactions among all DNA segment pairs. These maps show a complex cross-scale organization spanning megabase-pair compartments to short-ranged DNA loops. To better understand the organizing principles, several groups analyzed Hi-C data assuming a Russian-doll-like nested hierarchy where DNA regions of similar sizes merge into larger and larger structures. Apart from being a simple and appealing description, this model explains, e.g., the omnipresent chequerboard pattern seen in Hi-C maps, known as A/B compartments, and foreshadows the co-localization of some functionally similar DNA regions. However, while successful, this model is incompatible with the two competing mechanisms that seem to shape a significant part of the chromosomes’ 3D organization: loop extrusion and phase separation. This paper aims to map out the chromosome’s actual folding hierarchy from empirical data. To this end, we take advantage of Hi-C experiments and treat the measured DNA-DNA interactions as a weighted network. From such a network, we extract 3D communities using the generalized Louvain algorithm. This algorithm has a resolution parameter that allows us to scan seamlessly through the community size spectrum, from A/B compartments to topologically associated domains (TADs). By constructing a hierarchical tree connecting these communities, we find that chromosomes are more complex than a perfect hierarchy. Analyzing how communities nest relative to a simple folding model, we found that chromosomes exhibit a significant portion of nested and non-nested community pairs alongside considerable randomness. In addition, by examining nesting and chromatin types, we discovered that nested parts are often associated with active chromatin. These results highlight that cross-scale relationships will be essential components in models aiming to reach a deep understanding of the causal mechanisms of chromosome folding.

## I. INTRODUCTION

Mammalian genomes fold into a network of 3D structures that facilitate and regulate genetic processes such as transcription, DNA repair, and epigenetics^1;2^. Most discoveries derive from chromosome capture methods, such as Hi-C, which measure the number of contacts between DNA segment pairs and allow researchers to construct genome-wide 3D contact maps^3–5^. These maps show that chromosomes comprise a spectrum of 3D structures spanning a range of scales: megabase-scale A/B compartments, sub-megabase-scale Topologically Associated Domains (TADs), and short-ranged loops. Some of these structures are associated with epigenetic marks, active genes, and architectural proteins that reshape chromatin, such as CCCTC-binding factors (CTCF), cohesin complexes, and CP190^6–9^.

At first glance, Hi-C maps appear hierarchical, where DNA regions sharing high contact counts fold into larger and larger structures. This scheme is appealing because it proposes a simple folding mechanism leading to densely packed DNA without over-entanglement. It also predicts the existence of alternating megabase-sized 3D structures appearing in most Hi-C maps as plaid patterns^3;6;10^. More specifically, TADs tend to aggregate into sub-compartments (denoted A1, A2, B1, …, B4)^11^.

This folding scheme also posits that chromosomes form a perfect hierarchy. In other words, once two DNA regions join, such as two TADs, they remain in the same super-structure throughout the upstream folding hierarchy. This idea is the keystone assumption in several studie^s12-15^. While it can explain how A/B compartments form and foreshadows the co-localization of some functionally similar DNA regions, critical observations question the basic idea.

First, 3D communities are not necessarily contiguous DNA segments^16^. Assembling such disconnected communities into larger and larger building blocks inevitably leads to a non-perfect hierarchy. Second, if the hierarchy is perfect, it suggests that similar folding mechanisms act across several scales. However, this conclusion is inconsistent with the competing mechanisms that seem to form TADs and A/B compartments: loop extrusion and phase-separation^17;18^. Third, in a recent paper^19^, researchers fit a Gaussian polymer model to Hi-C data and recovered several established sub-structures—TADs, subTADs, A/B compartments, etc.—showing that they were not perfectly hierarchical.

This paper aims to unveil the actual folding by charting the cross-scale structural relationships from empirical data. In particular, we use data from Hi-C experiments that we recast into a weighted network of 3D interactions and use tools from network science to find the optimal community assembly while scanning through the networks’ layers of organization. By mapping the hierarchical relationships between these assemblies, we find that some nest, others segregate, and still others are not significantly different from random. To better understand these results, we propose a minimal folding model mixing perfect and random nesting. We also relate community nesting to established chromatin states. We discovered that communities associated with active transcription are more distinct and show significant nesting relative to the chromosome-wide average.

## II. MATERIALS AND METHODS

### A. Hi-C data treatment

We used Hi-C data for the human cell line GM12878 (B-lymphoblastoid)^4^, downloaded from the GEO database^20^. We used the MAPQG0 data set at 100 kilobase-pair (kb) resolution. Stored in matrix form, the Hi-C data contains the pairwise contact counts between DNA loci. We omit inter-chromosome contacts due to their low signal-to-noise ratio^21;22^.

We treat the Hi-C data as a DNA contact network. Each network node represents a 100 kb DNA segment, and the link weights are proportional to the number of measured Hi-C contacts. We use network methods (the generalized Louvain method, see Sec. II.B) to extract communities harbouring densely connected nodes that maintain fewer contacts with the rest of the network.

Before investigating the community structure, we normalized the raw Hi-C counts to reduce biases and to make fair comparisons between chromosomes that may vary in size up to one order of magnitude. In particular, we use the Knight-Ruiz (KR) matrix balancing^23^ implemented in gcMapExplorer^24^.

### B. The GenLouvain algorithm—detecting 3D communities in Hi-C data

To find communities in the Hi-C contact network, we use the generalized Louvain method (GenLouvain)^25;26^. It is a community detection method that takes advantage of so-called modularity maximization to find the optimal community division of a network. By “optimal,” we mean the community assembly that maximizes the number of internal contacts, measured by a so-called modularity function, with respect to a null hypothesis. A common null model is random rewiring keeping the node degree fixed. GenLouvain is a greedy optimization algorithm that starts with single-node communities and then searches for the optimal solution by generating trial node agglomerates and evaluating the modularity function.

We chose GenLouvain based on principled and practical aspects. First, it is a generalized version of one of the most intensively tested algorithms: Louvain^25^. Second, for practicality, the developer’s open-source codes are written in `MATLAB`, allowing us to modify essential parts such as the null-model term, which we will elaborate on in the forthcoming paragraphs.

One critical feature of GenLouvain is its resolution (or scale) parameter *γ*. This parameter allows us to sweep through the scales of the network and probe the network’s community spectrum. Furthermore, this parameter is closely related to the parameters capturing the relative tendency of intra-versus inter-group connections in the context of the stochastic block model^27^. Mathematically, *γ* is a part of the modularity function 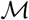, defined as

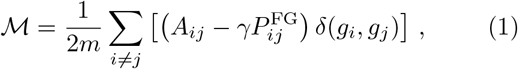

where *A_ij_* is the network’s adjacency matrix representing the weight of the edge connecting nodes *i* and *j*, and the summand is counted only if *i* and *j* belong to the same community, thus the Kronecker delta *δ*(*g_i_*,*g_j_*). In our case, *A_ij_* corresponds to the KR-normalized Hi-C matrix.

Furthermore, the second term of the summand in Eq. (1) represents our null hypothesis for the network’s “background” connectivity. Building on previous work^16^, we use the so-called fractal-globule (FG) null-model term 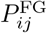 that assumes that the average interaction strength between two DNA segments, *i* and *j*, decays as a powerlaw with exponent −1. The FG null model term is

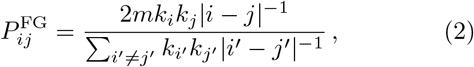

where the strength *k_i_* represents the sum of weights around node *i*, 2*m* = ∑_*i*_ *k_i_* is a normalization constant, and ∝ 1/|*i* — *j*| is the expected amount of reduced interaction as a function of the one-dimensional distance separating nodes *i* and *j*. This decay follows the fractal-globule scaling^16;28;29^ that agrees with chromatin contact decay in Hi-C experiments, where the exponent is approximately −1.08^3^. Note that we may use any decay exponent in Eq. (2). For example, −0.75 matches better with within-TAD contacts,^30^, but we kept −1 as our 3D communities are larger than TADs.

### C. Network nestedness

By varying the resolution parameter *γ* embedded in the GenLouvain algorithm, we scan through the scales of chromosomes’ 3D organization. While scanning, we keep track of uninterrupted DNA segments—we will refer to these segments as *domains* in Sec. III—and how they distribute between the communities as the scale changes. This allows us to chart cross-scale folding relationships.

To better understand these relationships and quantify the deviations from a perfect hierarchy, we use an approach developed for ecological networks^31;32^. Designed for interacting species pairs, say plants and pollinators, this approach rests on a nestedness metric, called *N_ij_*, measuring how many plants two pollinators have in common compared to a random benchmark. In our case, we track how many DNA segments (domains) two 3D communities share, given that they appear at different hierarchical levels (different *γ* values)—by construction, two communities at the same hierarchical level do not share any domains. We illustrate the philosophy behind *N_ij_* in Fig. 1.

**FIG. 1.**
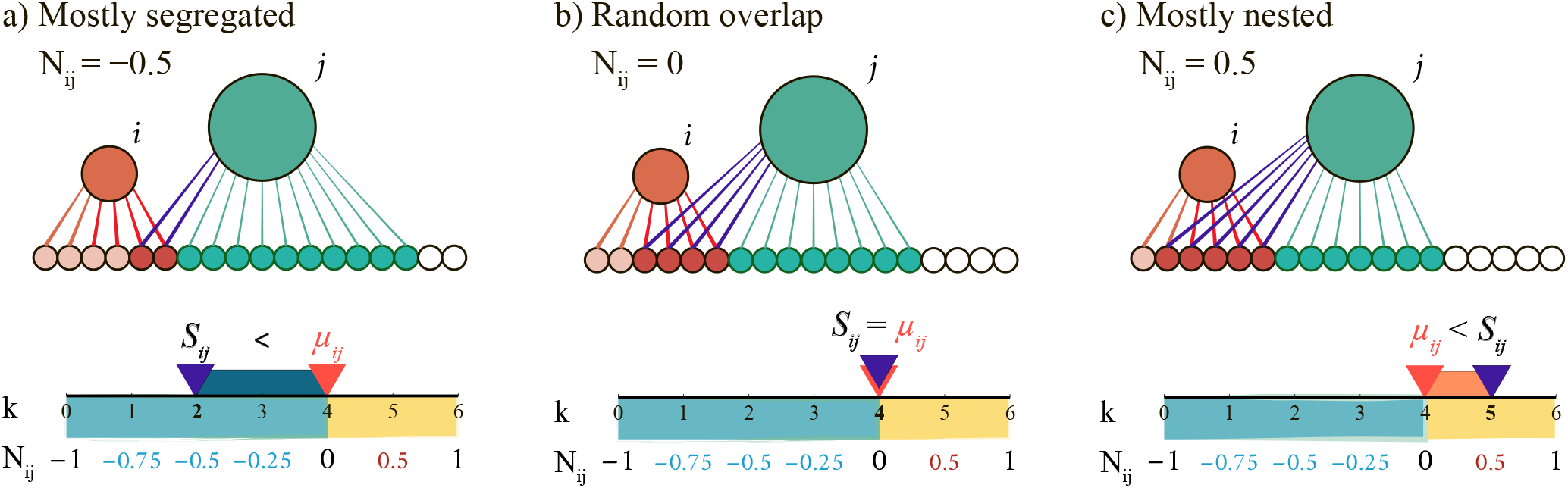
Three examples of nestedness (*N_ij_*) in a simple bipartite network. The networks in panels (a)–(c) have the same number of nodes in each layer—18 domains (small circles) and two communities (*i* and *j*, large circles)—and the same number of links (18) but connected differently to achieve varying nestedness. Below each network, we illustrate how we calculate *N_ij_* using Eqs. (4)–(7). On the horizontal *k*-axis, we indicate the number of shared nodes *S_ij_* for the community pair and the expected overlap *μ_ij_* calculated from Eq. (3). The panels (a)–(c) show three essential *N_ij_* regimes (*μ_ij_* = 4 for all of the cases). (a) Mostly segregated (*S_ij_* < *μ_ij_*, *N_ij_* = −0.5). Because *S_ij_* = 2 and *μ_ij_* = 4, the *i* and *j* communities are half-way to full segregation. We illustrate this with a dark-blue stripe covering half the 0 ≤ *k* ≤ 4 = *μ_ij_* range. (b) Random overlap (*S_ij_* = *μ_ij_*, *N_ij_* = 0). The number of shared nodes equals the random expectation. (c) Mostly nested (*S_ij_* > *μ_ij_*, *N_ij_* = 0.5). Here *i* and *j* share one domain more than expected (*S_ij_* = 5). This yields *N_ij_* = 0.5 because their overlap is at the midpoint between the random and the maximum overlap (*S_ij_* = 6) that would result in ideal nesting (*N_ij_* = 1). We illustrate this with the orange stripe spanning half of the range *μ_ij_* (= 4) ≤ *k* ≤ 6. This example shows that *N_ij_* measures the relative overlap compared to what is achievable given the link density rather than absolute numbers.

The *N_ij_* metric benchmarks the community overlaps to a combinatorial link redistribution, normalized to vary between −1 and +1. These endpoints indicate complete network segregation (*N_ij_* = −1) or perfect nesting (*N_ij_* = +1). When perfectly nested, the larger community engulfs the domains in the smaller ones. When completely segregated, the communities do not share any domains^1^. In a perfect hierarchy, like a phylogenetic tree, the nestedness is either +1 or −1, indicating full nesting or complete segregation. But in a more complex multiscale structure, *N_ij_* takes any value between these two extremes because the communities may share more or fewer domains relative to a random overlap. We note that *N_ij_* is normalized so that the midpoint *N_ij_* = 0 represents random overlap and that *N_ij_* = ±*x* indicates the same relative proportion *x* of segregation or nesting. We exemplify this property in Fig. 1 using a small bipartite network having varying nestedness: (a) mostly segregated (*N_ij_* = −0.5), (b) random overlap (*N_ij_* = 0), and (c) mostly nested (*N_ij_* = 0.5).

We go through several steps to calculate *N_ij_*. First, we extract the overlap *S_ij_* between two communities *i* and *j* from data—we study nestedness in empirical (Hi-C-derived) and simulated data. Second, we calculate the expected overlap *μ_ij_* assuming a random arrangement. Denoting *d_i_* as community *i*’s internal number of domains, *μ_ij_* is^31^

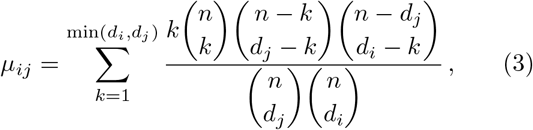

where *n* is the total and *k* is the shared number of domains. Next, we shift *S_ij_* by *μ_ij_* to center the expected overlap for random arrangement at zero and normalize so that *N_ij_* ∈ [−1, 1]:

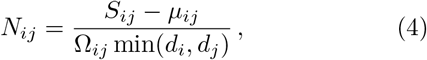

where *Ω_ij_* is the maximum or minimum achievable overlap, depending on if *S_ij_* > *μ_ij_* or *S_ij_* < *μ_ij_*. In these cases, we calculate *Ω_ij_* as

a. *S_ij_* > *μ_ij_*

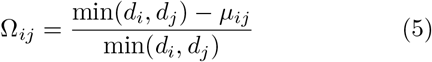
b. *S_ij_* < *μ_ij_*, which is further classified into two cases:

i. *d_i_* + *d_j_* — *n* < 0

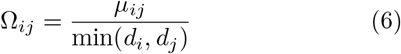
ii. *d_i_* + *d_j_* — *n* ≥ 0

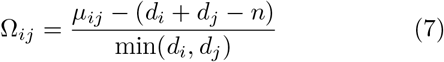

#### 1. Significant community overlap and *p*-values

In addition to expected the overlap *μ_ij_*, we calculate the likelihood that two communities share *S_ij_* domains given the random null hypothesis. Under this hypothesis, the probability that *S_ij_* = *k* is^31;32^

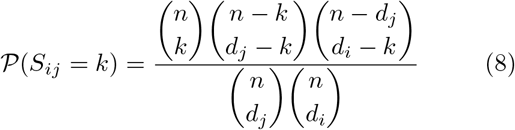

To assign *p*-values to the observations, we sum 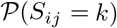 over *k*. However, depending on if *k* is smaller or larger than *S_ij_*, we must separate two cases

i. *k*_obs_. ≤ *S_ij_*,

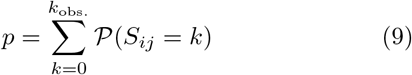
ii. *k*_obs_. ≥ *S_ij_*

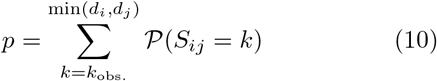

In our analyses, we set the significance threshold to *p* ≤ 0.025 to distinguish significant from random overlap.

### D. Chromatin states and folds of enrichment

In Sec. III.C, we study cross-scale nestedness among communities associated with specific chromatin states. To calculate chromatin enrichment, we used published data integrating several resources (e.g., ChIP-seq and RNA-seq) to partition the genome into 15 chromatin types.^33^ Derived from a multivariate Hidden Markov Model (HMM), these states are (S1–S15): Active Promoter (S1), Weak Promoter (S2), Inactive/poised Promoter (S3), Strong Enhancer (S4, S5), Weak/poised Enhancer (S6, S7), Insulator (S8), Transcriptional transition (S9), Transcriptional elongation (S10), Weakly transcribed (S11), Polycomb-repressed (S12), Heterochromatin (S13), and Repetitive/Copy number variation (S14, S15).

We downloaded the HMM data from ENCODE (human cell line GM12878)^34^. The data is a genome-wide list of start-and-stop coordinates for each HMM state, where each instance is called a “peak”. To determine the HMM content in a long DNA stretch, say a community, we count the number of peaks belonging to each of the 15 states. Because some HMM peaks may cross community borders, we count the number of peak starts.

Next, to calculate the enrichment, we use a hypergeometric test that benchmarks the HMM content in a community to the chromosome-wide random expectation (sampling without replacement). The test goes through the following three steps.

1. Get community content from HMM data. We denote the number of peaks for each state as *k_X_*, *X* = S1,…, S15.
2. Calculate the expected number of *X* peaks 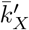 given the community’s total peak count *n* as 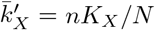, where *N* is the total number of peaks in the chromosome (including all HMM states), and *K_X_* is the number of *X* peaks in the chromosome.
3. Calculate the *p*-value for 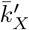 under the hypergeometric null hypothesis. If less than the significance threshold of 0.05, we consider the community enriched or depleted in HMM state *X* (twosided test). However, because we make multiple comparisons, one for each HMM state, we correct the *p*-value to reduce the false discovery rate. We do this using the Benjamini-Hochberg procedure^35^ implemented in Python statsmodels^36^. We set the false discovery rate to 0.05.

After going through all communities using this procedure, they get labeled as “enriched” or “depleted” in each of the 15 chromatin states. We point out that one community can be enriched in several HMM states.

In addition, to make our analysis more tractable when studying the nestedness of different chromatin types, we make a coarser classification and partition the communities into four large groups, A–D. These groups reflect the overall HMM state enrichment: A: Active promoters (S1–S2), B: Enhancers (S4–S7), C: Transcribed regions (S9–S11), and D: Heterochromatin (S3 and S12–S15).

## III. RESULTS

### A. Distant domains aggregate into 3D communities spanning a range of scales

To illustrate how the 3D communities partition the chromosome, we superimpose GenLouvain-derived communities as squares along the Hi-C map’s diagonal in Fig. 2(a). By assigning each community a unique color, we see that some 3D communities contain distant DNA segments. This community type—a distributed assembly of DNA segments—widens commonly used 3D partitions, like TADs, that assumes contiguous DNA stretches^37^.

**FIG. 2.**
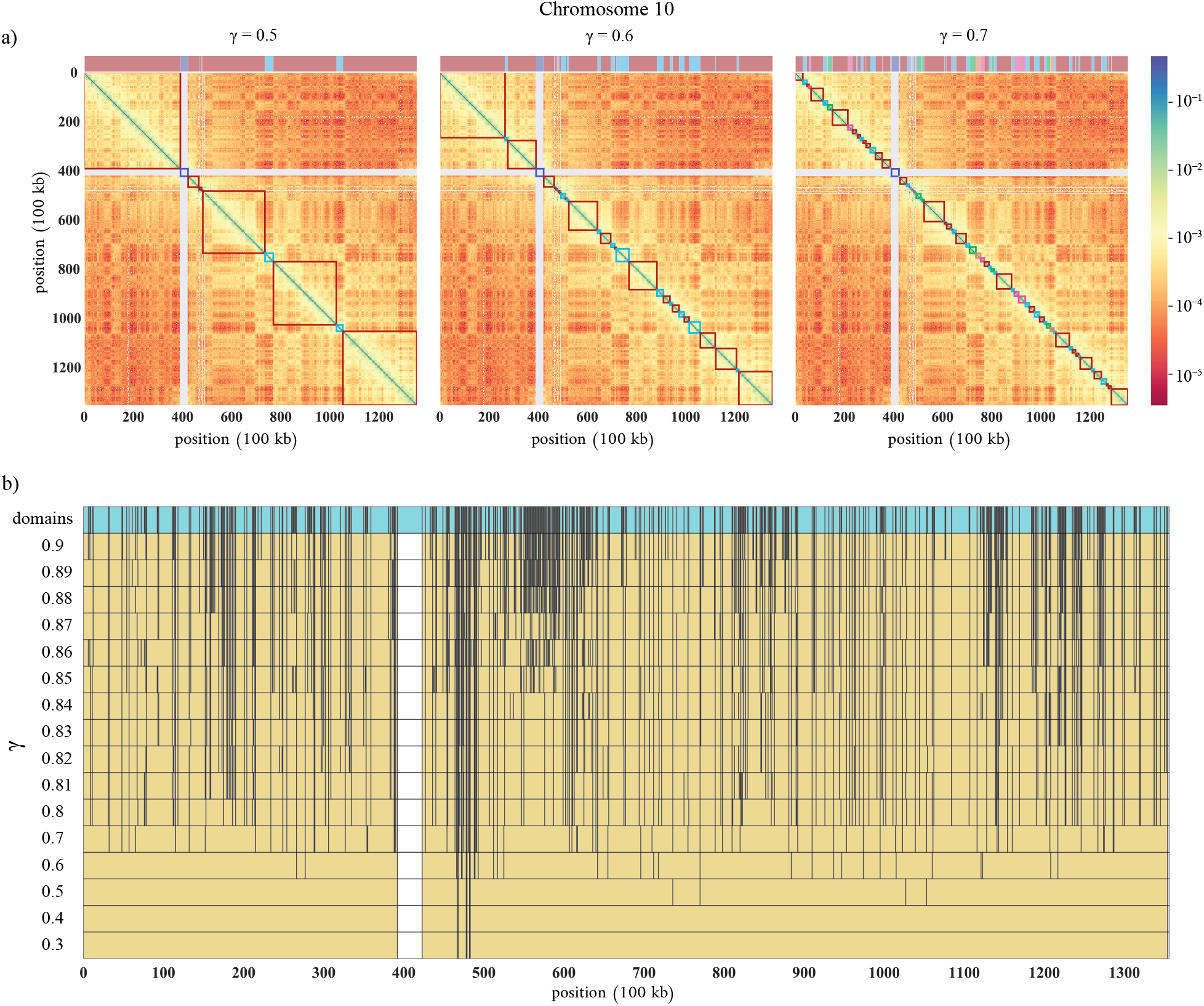
Hi-C maps, 3D communities, and domains. (a) Hi-C maps where the red-to-blue pixel colors are a proxy for short-to-long 3D distances. The squares decorating the map’s diagonals represent GenLouvain-derived 3D communities for three *γ* values (0.5, 0.6, and 0.7). Above each map, we show the community coverage as a colored stripe. Having unique colors, we observe that the communities comprise scattered linear DNA segments. The white cross shows the centromere. (b) Community borders and coverage across chromosome 10 for 16 *γ* values. The upper turquoise stripe shows DNA stretches that never split for 0 < *γ* ≾ 1. We refer to these indivisible regions as domains. The smallest 3D domain is 100 kb long, which is the resolution limit of the Hi-C data set we use.

Furthermore, Fig. 2(b) shows that some DNA stretches rarely break across a wide range of *γ*. We call these indivisible pieces “irreducible domains”. We collect them by we collecting borders of intact DNA segments across many *γ* values into one list. We show the domains in the upper turquoise stripe in Fig. 2(b) and their size distribution in Fig. S4 (Supplementary Material). Admittedly, making *γ* large enough, we break even the domains into smaller linear DNA pieces so that eventually every Hi-C bin (100 kb) represents one domain. However, we do not cover this extreme limit here.

### B. 3D communities do not form perfect hierarchies

Figure 2 suggests that 3D communities have complex cross-scale relationships. To better visualize such relations, we constructed a hierarchical tree from the same Hi-C data set (chromosome 10), showing how domains, the least divisible DNA regions, join into large 3D structures that, in turn, make up 3D communities (Fig. 3).

**FIG. 3.**
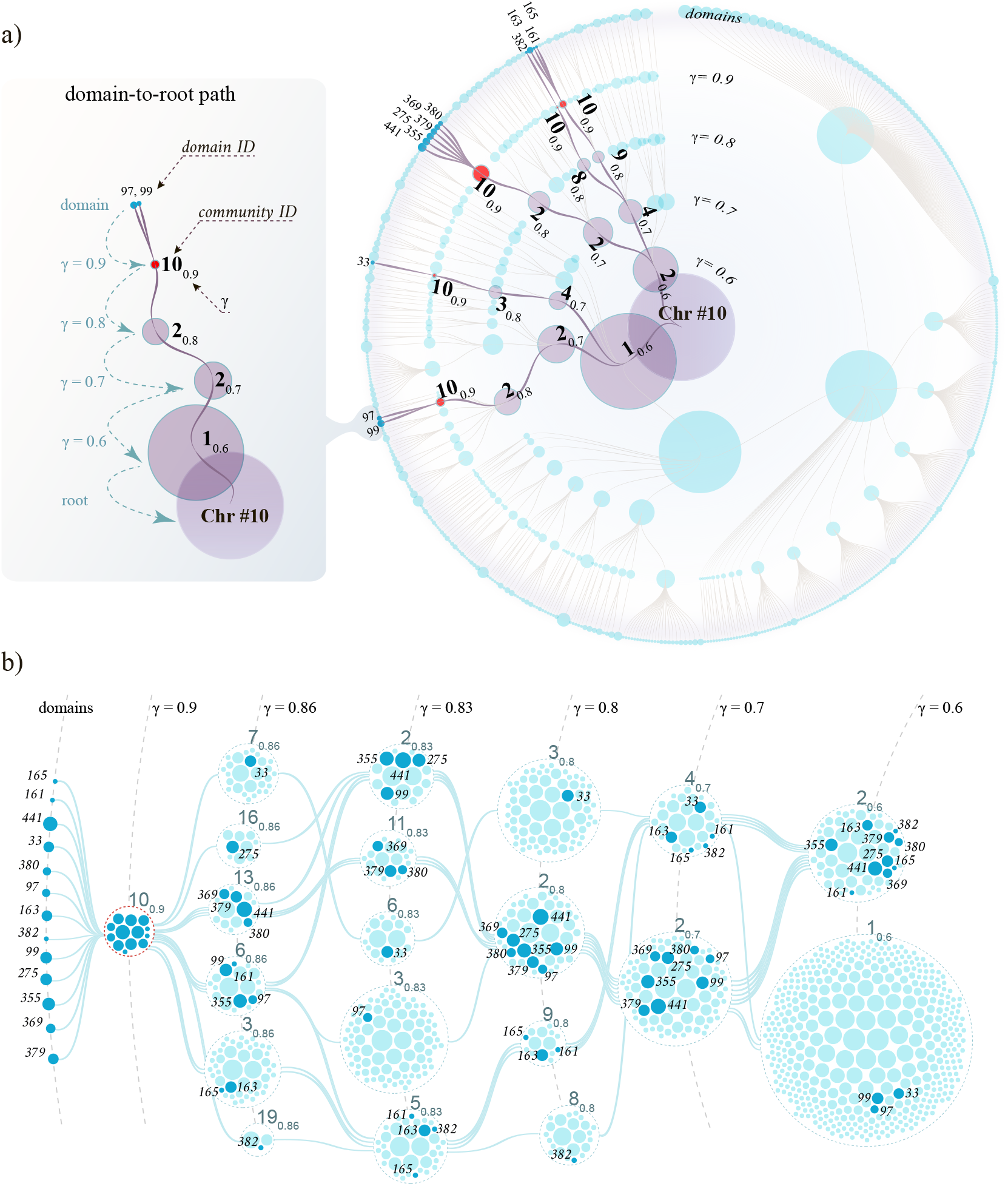
Cross-scale community organization in chromosome 10. (a) Circular tree showing how domains (filled circles on the outer rim) merge into larger and larger 3D structures (filled circles on the inner rings). Each ring represents one value of GenLouvain’s resolution parameter *γ*, and the diameters of the filled circles are associated with their DNA length (measured by the number of Hi-C bins). The red circles mark delocalized 3D structures forming a single 3D community at *γ* = 0.9 (denoted 10_0.9_). The dark links show folding trajectories for the domains passing through 10_0.9_ towards the root. The left panel shows a two-domain folding path and defines our label convention. We plotted the tree using RAW Graphs^38^. (b) Joining and splitting of the 13 domains belonging to the community 10_0.9_. These domains (filled dark-blue circles) pass through the 3D communities (open circles), joining other domains (filled light-blue circles). The edges connect 3D communities with dark-blue domains. We also highlighted these folding pathways in (a) (dark links).

To construct the tree in Fig. 3, we first extracted chromosome 10’s domain list and calculated the optimal community division associated with a few *γ* values. Next, we stored the folding pathways of all domains by tracing how their community memberships change with *γ* (Fig. 3(a), left). The circular tree illustrates the collection of all these pathways, where the links indicate how domains (filled circles on the outer rim) assemble into 3D structures (filled circles on the inner rings). Each ring corresponds to one *γ* value, i.e., one organization scale, and the filled circles’ diameter symbolizes their DNA content.

The tree in Fig. 3(a) looks hierarchical. But a more complex pattern emerges if factoring in the 3D communities. To this end, we print the community ID next to a few filled circles (black numbers). In addition to the ID, we add a subscript to indicate the *γ* value (e.g., ID*_γ_*). However, we omit ID numbers for the domains on the outer rim. Instead, their numbering denotes sequential ordering along the chromosome (e.g., domain 2 is next to domains 1 and 3, etc.).

Interestingly, the same community ID appears several times within one *γ* ring. One example is the community 10_0.9_, highlighted in red, that appears five times on the *γ* = 0.9 ring. This community contains 13 domains scattered over the chromosome, as seen from their non-consecutive ID numbers. But even if scattered, they belong to the same 3D community that is a part of the optimal network division (according to GenLouvain). Delocalized domains forming communities this way is a hallmark of imperfect hierarchical folding.

To further exemplify this observation, we depict the folding pathways of the 13 individual domains belonging to the community 10_0.9_ in Fig. 3(b). By following the folding paths (edges) from left to right, we see that these domains (filled dark-blue circles) start in the same community and then split apart to become members of other 3D communities having different domain content (light-blue circles). Going even further to the right, 10 out of 13 dark-blue domains join yet again into one huge community (at *γ* ≾ 0.6). Again, this complex merging- and-splitting behavior is far from a perfect hierarchy. For clarity, we highlighted community 10_0.9_’s folding pathways as dark lines in (a) connecting the violet circles.

### C. Quantifying chromosome nestedness

Figure 3 shows that domains mix between 3D communities as they approach the tree’s root. This finding suggests that the folding mechanics is not perfectly hierarchical. To quantify deviations from being perfect, we calculate the pairwise community-domain overlap relative to random chance between two communities, *i* and *j*, belonging to different tree rings. To this end, we use a normalized nestedness metric, denoted *N_ij_*, that varies from −1 to +1. These two extreme points indicate complete segregation (*N_ij_* = −1) and perfect nesting (*N_ij_* = 1). When *N_ij_* = 0, the overlap is not different from being random. We outline the explicit calculations and some of *N_ij_*’s critical properties in Sec. II.C and show a schematic in Fig. 4(a).

**FIG. 4.**
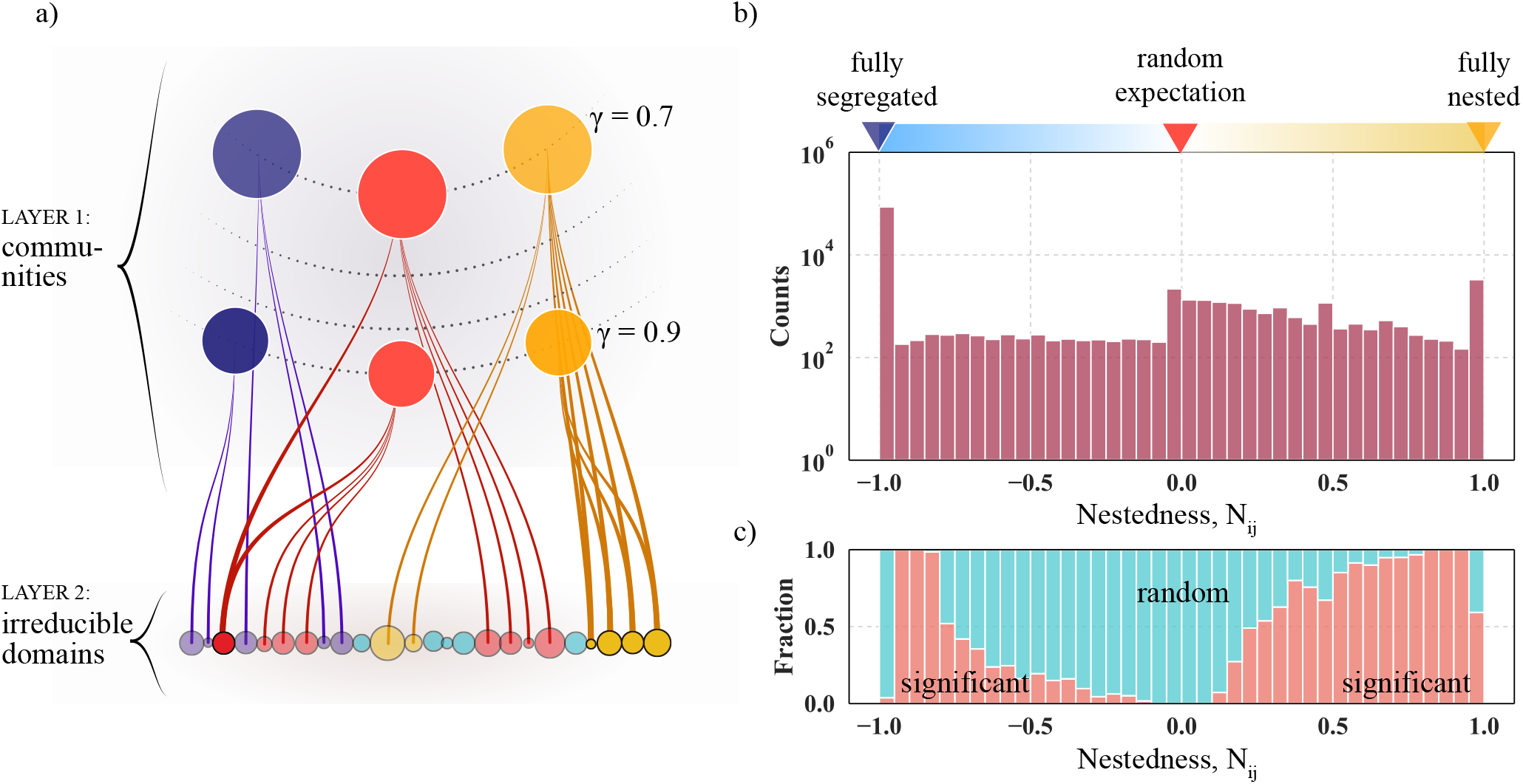
Nestedness of community pairs in human chromosomes 3, 5, 10, and 22. (a) Schematic community-domain overlap in three cases: fully segregated (*N_ij_* = −1, violet), random (*N_ij_* = 0, red), and fully nested (*N_ij_* = 1, yellow). Layer 1 contains communities belonging to different *γ* values (dotted lines). The bottom layer shows the irreducible domains, and the edges indicate community memberships. (b) Nestedness histogram (Nj) for chromosomes 3, 5, 10, and 22. For each chromosome, we derived communities from 16 *γ* values. The peaks at ±1 suggest that several communities are segregated (−1) and nested (+1). However, there also exist significant intermediate levels of nestedness. The stripe overlaying the histogram indicates what we classify as fully segregated or nested according to the nestedness metric outlined in Sec. II.C. We show the nestedness of individual chromosomes in Fig. S3, and we visualize *N_ij_* distributions for individual *γ*-pairs in Supplementary Material, Fig. S5 (chromosome 10). (c) Significant versus random community nestedness. As outlined in Sec. II.C, we filter community overlaps having *p*-values ≤ 0.025 and show the relative proportions of significant and random overlap associated with the *N_ij_* histogram in (a). The colors indicate significant (orange) and random overlaps (blue-green).

To study the cross-scale nestedness in Hi-C-derived trees, like Fig. 3, we calculated *N_ij_* across several *γ* values in four chromosomes (3, 5, 10, and 22); we choose these to mix large, intermediate, and small chromosomes. Plotting the *N_ij_* histogram for all chromosomes in one graph, we find that the distribution has two pronounced peaks at ±1 and a flat but slightly right-skewed intermediate region [Fig. 4(b)]. These two peaks indicate that some communities segregate (−1) while others nest (+1), just like in a perfect hierarchy that is either completely segregated or fully nested. However, the histogram’s intermediate *N_ij_* region is not zero and thus differs from an ideal hierarchy. This telltales that the 3D folding blends hierarchy-breaking contacts where some are possibly random.

To separate significant from random overlaps in Fig. 4(b), we calculated the probability that two 3D communities, having sizes *d_i_* and *d_j_*, share *k* domains in a random assignment—we defer all details to Sec. II.C. Based on this probability, we associate *p*-values to each *N_ij_* observation. Setting the threshold to *p* ≤ 0.025, we count the fraction of significant observations and illustrate the relative proportions in Fig. 4(c). In orange, we highlight significant overlaps. In blue-green, we indicate overlaps that are indistinguishable from being random. From panel (c), we make three key observations. First, the most segregated part (−1) is almost entirely blue-green and thus classified as insignificant. Second, roughly half of the perfectly nested communities (+1) share significant domain overlaps. Third, two regions show substantial overlaps that err on the side of segregation (−0.95 < *N_ij_* < −0.8) and nesting (0.65 < *N_ij_* < 0.95). The remaining data points appear random, particularly those surrounding *N_ij_* = 0.

From Figs. 2–4, we conclude that chromosomes fold into complex hierarchies that mix nested and segregated parts. On average, however, the nesting is close to being random 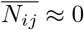 if neglecting the −1 peak that skews the average (if included, it is 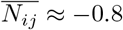). But since the distribution is so broad, community pairs show substantial differences where some are completely segregated, others are perfectly nested, and the rest is somewhere in between. This finding sheds new light on the hierarchical chromosome paradigm underlying several papers. Finally, in Fig. S5, we show community overlaps between specific *γ* pairs.

### D. Modeling non-nested chromosome folding

Several papers assume that linear DNA regions, like TADs, form higher-order structures by folding into each other in a perfect hierarchy (e.g.,^12–15^). However, our data show that the nesting is more complex (Figs. 2–4). To better understand this disconnect, we propose a model for semi-nested chromosome folding. At the core, the model assembles ideally nested domain groups, consistent with the significant nestedness seen in the *N_ij_* histograms (Fig. 4). Then, we break this pattern by reshuffling some domains among the communities. We denote the critical reshuffling parameter *Q* that represents the probability that two domains will change community memberships. Below, we outline the *Q* = 0 (perfect hierarchy) and *Q* > 0 limits separately.

#### 1. Perfect hierarchical folding (*Q* = 0)

To achieve ideal hierarchical folding, we agglomerate domains into superstructures and superstructures into yet larger superstructures, following a few simple steps. First, we calculate the pairwise domain-domain interaction strength from their average Hi-C contact frequency (domains typically consist of several Hi-C 100 kb bins). Second, we select the domain pair having the strongest interaction and merge them into a superstructure. Then we replace the two merged domains in the list of pairwise interactions with the new superstructure and join the next most interacting pair. Regardless of choice, this scheme yields a new superstructure at each iteration. Notably, the algorithm does not only merge linearly adjacent domains.

Once we merge all domains into a giant superstructure, we use the domains’ folding paths to organize the super-structures into a circular tree [Fig. 5(a)]. However, unlike the Hi-C derived tree in Fig. 3, the rings in Fig. 5(a) do not represent different *γ* values. Instead, they show consecutive mergers of the superstructures. Because some branches are so deep (> 10 steps), we show only the last five merging events and put all the domains on the outer (sixth) rim.

**FIG. 5.**
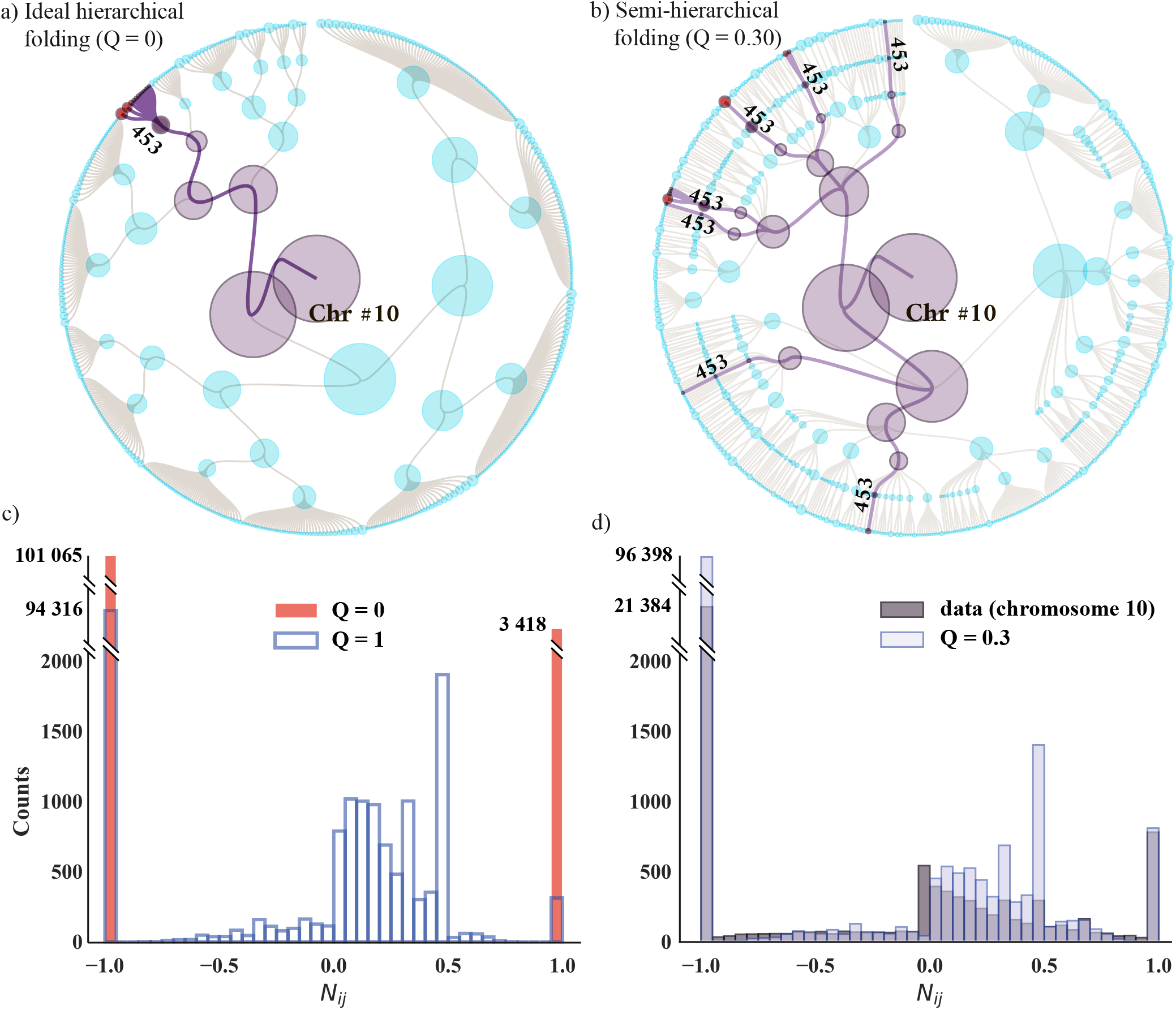
Hierarchical and semi-hierarchical models of chromatin folding for human chromosome 10. (a) Ideal hierarchical folding (*Q* = 0). Filled circles on the outer rim represent domains; the root symbolizes the entire chromosome. We align the domain aggregates (superstructures) with the inner tree rings, each defining a scale of organization. We select a few domains (red-filled circles) and show their domain-to-root paths with thick edges. These domains assemble into yet larger structures (violet) at every inner ring. As soon as the domains merge into a superstructure labeled ‘453’ (dark violet), they never split apart. (b) Semi-hierarchical folding (*Q* = 0.30). As in (a), we color the domains in red that merge into a superstructure ‘453’ and highlight their folding paths with thick edges going from the outer rim to the root. Unlike (a), node ‘453’ is scattered across seven tree branches. Thus, ‘453’ only partially nests into larger structures and the domains split and reunite when approaching the root. (c) Nestedness histogram when *Q* = 0 (ideal hierarchy, red bars) and *Q* = 1 (random nesting, open bars). When *Q* = 0, we see two peaks at *N_ij_* ± 1, indicating complete segregation and full nestedness. When *Q* = 1, the domains are fully randomized between the superstructures. While there is still perfect nesting and segregation (as we expect from the random null hypothesis in Sec. II.C), there is also partial overlap for −0.8 < *N_ij_* < 0.8. (d) Nestedness histogram with some randomness (*Q* = 0.30, light-blue bars) overlaying the actual GenLouvain-derived data for chromosome 10 (dark-grey bars). We produced (a) and (b) using RAW Graphs^38^.

To illustrate that this scheme produces an ideal hierarchy, we select a group of 12 domains (red-filled circles, outer rim) and highlight their folding paths across the tree with heavy links. After forming one super structure (‘453’, dark violet), this domain group stays intact as it joins more and more domains forming increasingly larger superstructures (violet circles). This exemplifies that domains never split apart once they end up in the same superstructure. However, this behaviour contrasts with what we observed in Fig. 3, where domains split and merge as they form communities. Therefore, this simple description cannot explain actual chromosome folding. We point out that the *Q* = 0 limit is nearly identical to the so-called metaTAD algorithm^12^.

#### 2. Hierarchical folding with randomness (*Q* > 0)

The model yields a perfect domain hierarchy when the reshuffling parameter *Q* is zero. However, the actual folding patterns appear more complex. We exemplified this in Fig. 2(b), illustrating the cross-scale folding paths of 12 domains in chromosome 10. Following these paths, we note they do not perfectly correlate as they would in a perfect hierarchy. While some domains often stay together, others split only to reunite later. This represents the feature we aim to mimWhenle a fraction of domains between the communities, restricting the reshuffling to communities within the same level of organization. The number of domains we interchange is proportional to *Q*. Algorithmically, we follow these three steps. (1) Go through all superstructures in the same organizational level [one ring in Fig. 5(a)] and identify the domain IDs and superstructure memberships. We exclude domains that do not yet belong to any community. (2) Select two of these domains randomly and swap their superstructure memberships with probability *Q*. (3) Repeat (1)–(2) until we exhaust all domain pairs, excluding those we already interchanged. If one domain remains without a pair, we keep its superstructure membership. Next, we pick another tree ring and repeat steps (1)–(3).

By varying the parameter *Q*, we retrieved several *N_ij_* distributions. To find the optimal *Q*_opt_. —the *Q* that produces the *N_ij_* distribution that is most similar to the real data—we utilized a Kolmogorov-Smirnov test. This test gave *Q*_opt_. ≈ 0.3 (Supplementary Material, Fig. S8). Fig. 5(b) shows the associated domain folding paths.

In contrast to *Q* = 0 in Fig. 5(a), Fig. 5(b) shows that the hierarchy breaks when *Q* > 0. We highlighted the domains forming the same superstructure we studied in (a) (‘453’, dark violet) to better see the difference. Like in (a), this superstructure has 12 domains (11 out of 12 are the same). But unlike (a), superstructure 453 appears in different tree branches. This better reassembles the Hi-C derived tree in Fig. 3(b), where domains merge that do not have identical domain-to-root folding paths.

We point out that the tree’s backbone formed in this way is identical to the *Q* = 0 case, but the domain memberships differ. Therefore, we foreshadow that this model is valid for small *Q*. But as we show in the following section, this is enough to reproduce the actual nestedness distribution in Fig. 4.

#### 3. Nestedness for hierarchical and semi-hierarchical folding

To study how the *Q* parameter in the model affects superstructure nestedness, we calculated and studied the *N_ij_* histograms. Just as in Sec. III.C, we calculate these histograms by going through all superstructure pairs, omitting those belonging to the same tree ring, and counting the number of shared domains (Sec. II.C). We show three cases in Figs. 5(c) and (d): perfect hierarchy (*Q* = 0), full randomness (*Q* = 1), and intermediate randomness (*Q* = 0.30). All three cases build on domains derived from chromosome 10. Below, we discuss each case separately.

As expected for the ideal hierarchy, Fig. 5(c) has two isolated peaks at ±1 (red bars), indicating that the communities are either fully nested or fully segregated. Put differently, the structure is “modular.” These peaks also appear in the complete randomness limit (*Q* = 1). However, the +1 bar is lower relative to the *Q* = 0 case, and there is a distribution of *N_ij_* values surrounding *N_ij_* = 0, albeit not as wide as the actual data. We interpret this as the domain reshuffling split several nested communities while keeping the segregation primarily intact.

To better mimic the real data, we tweaked *Q* to reassemble the actual *N_ij_* histogram. In panel (d), we show the *Q* = 0.30 case overlaying the empirical data for chromosome 10. Apart from underestimating the histogram for negative *N_ij_* values and overestimating it for large values, the two histogram lies on top of each other for the most part. This shows that the reshuffling parameter must not be large for the model to reproduce the nestedness data in Fig. 4(b). About 30% domain redistribution seems enough.

### E. Nestedness and chromatin states

In Fig. 4, we found that some communities nest and others segregate. Also, Fig. 5 showed that we could reproduce the chromosome-wide nestedness distribution by slightly breaking an otherwise perfect folding hierarchy. This section analyzes if this behavior is associated with specific chromatin types.

To this end, we take advantage of published data that partition the genome into 15 chromatin states^33^. However, to make the analysis more tractable, we aggregate these states into four groups A–D, and study their pairwise nestedness. The groups are: promoters (A), enhancers (B), transcribed regions (C), and heterochromatin (D) (see Sec. II.D for complete definitions). To assign communities to these groups, we calculate folds of enrichment for each of the 15 chromatin states relative to the chromosome-wide average. We then use the hypergeometric statistical test to judge the enrichment significance (see Sec. II.D for details). Notably, because one community may enrich several chromatin states, it can belong to several A–D categories.

Next, we go through all community pairs to extract their nestedness *N_ij_* and chromatin group (A–D). Then we plot *N_ij_* histograms for all paired combinations—AA, AB, AC, AD, BB, etc. We show these histograms as panels in Fig. 6(b), where the light blue background portrays the entire chromosome’s nestedness (we use data from chromosome 10). The diagonal panels represent community pairs having the same chromatin type (AA, BB, CC, and DD). These pairs nest more than the rest of the chromosome as the *N_ij_* distributions skew to the right. This observation differs from DD, which seems to follow the chromosome’s overall nestedness distribution. To lend quantitative support to these observations, we performed a Kolmogorov-Smirnov test, which compares the cumulative distribution functions of the histograms AA, BB, CC, and DD (Supplementary Material, Fig. S6).

**FIG. 6.**
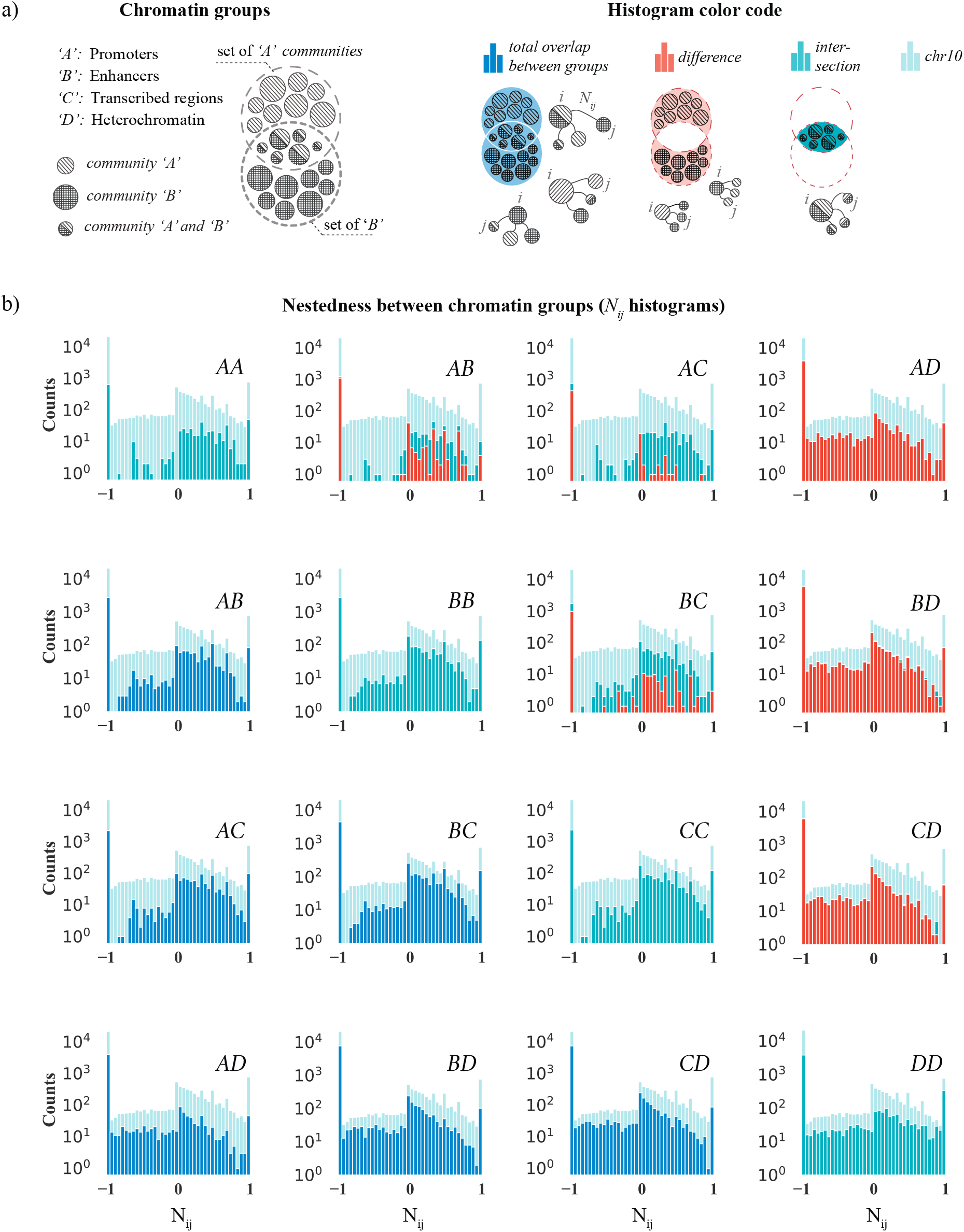
Chromatin type and cross-scale nestedness between community pairs in chromosome 10. (a, left) Each chromatin cross pair (AB, AC, AD, etc.) has three community types. For example, these may enrich A (“Promoters”), B (“Enhancers”), or A and B. The large dashed circles represent the Venn diagram of all A or B community types (small filled circles). The set illustrates that one community can be in one of three categories: enriched with A (upper), B (lower), or both (intersection). (a, right) Schematic illustrating the color codes used in the nestedness histograms associated: complete overlap (dark blue), the difference (dark red), and the intersection (light blue). In pale blue, we indicate the chromosome-wide average of chromosome 10. (b) Nestedness distributions (*N_ij_*) for 10 combinations of chromatin types A–D. The diagonal panels show the nestedness histograms for community pairs belonging to the same chromatin type (AA, BB, etc.). The off-diagonal panels show the other six paired combinations (AB, AC, AD, etc.); see panel (a) for detailed descriptions. The faint pale blue background in all histograms portrays the complete nestedness histogram from chromosome 10 (like Fig. 4).

However, we could argue that the folding structure is segregated rather than nested because the average *N_ij_* is negative in all diagonal panels; it becomes negative due to the large peak at −1 skewing the average. But as we showed in Fig. 4, this peak represents mostly random segregation (admittedly, roughly half of the +1 peak is also random), and the significant overlaps mostly appear for *N_ij_* > 0 where AA–CC histograms carry heavy weight. Therefore, we conclude that these chromatin groups nest more than the chromosome average and that the nesting is significant.

Furthermore, the off-diagonal panels in (b) show the *N_ij_* histograms for the six cross pairs, AB, AC, etc. But as noted above, some communities may enrich two groups simultaneously, say A and B. So when studying the AB cross-pair, it is natural to analyze separately communities enriched in A, B, or both, those enriched in A and B simultaneously, or those enriched in only A or only B. We depict these combinations and the color coding in panel (a), where the large dashed circles encompass all communities flagged as A or B. At the circles’ intersection, communities are enriched in A and B (half-filled circles).

The blue histograms in the lower triangular part of the panel (b) show the nestedness among communities belonging to the broadest class (e.g., A, B, or both). These off-diagonal histograms show that A, B, and C types tend to nest with each other (panels AB, AC, and BC), similar to AA–CC along the diagonal. In contrast, their overlap with D shows a wider variability reassembling the chromosome-wide *N_ij_* distribution, apart from the dip close to *N_ij_* = 1, hinting that A–C nest less with D than is expected. This observation likely reflects that A–C broadly belongs to what is commonly referred to as “active chromatin” and D is “inactive chromatin” (e.g., measured by low or high RNA expression levels). In addition, a more granular study analyzing all 15 chromatin states showed that the five states making up group D rarely enrich more than random alongside the others in A, B, and C. This differs from the A–C communities, where the internal chromatin states often co-appear. This explains why the significant nesting with group D is relatively scarce (see AD, BD, and CD panels).

In the upper triangular part in panel (b), we show stacked *N_ij_* histograms for the other two more restricted cross pairs (e.g., communities simultaneously enriched in A and B, or only A or B). In blue-green, we represent the intersection (e.g., A and B), and orange symbolizes the difference (e.g., only A or only B). Admitting that the sample size is relatively small, we note that the AB, AC, and BC histograms have a more significant fraction of data points for positive *N_ij_* values than those in the lower triangle, indicating more nesting. However, the AD, BD, and CD histograms remained almost identical.

In summary, when studying the cross-scale community nestedness, our data suggest that 3D communities belonging to “active chromatin” tend to nest more than the chromosome-wide average and appear to segregate from “inactive chromatin.” The data also indicates that communities embedded in inactive chromatin seem to have substantial random cross-scale overlaps.

### F. Active chromatin appears more hierarchical than inactive

To better understand the implications of the results in Fig. 6 regarding the chromosome’s 3D organization, we quantified how well different 3D communities partition the Hi-C network and if solid or weak divisions are associated with the chromatin groups A–D. To this end, we calculated the modularity associated with the GenLouvain-derived communities 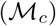. To calculate 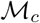, we use Eq. (1) and sum only those terms belonging to the same community. If the modularity scores high, the internal nodes interconnect more than the background. If scoring low, they connect less (See Sec. B for an explanation). We recover the global modularity 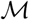 in Eq. (1) by summing over all communities, 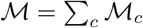.

The community modularity varies significantly within and between chromatin groups A–D (Fig. S9). We also found that 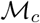 grows linearly with the community sizes (number of domains) (Fig. S9). Therefore, to make a fair comparison, we plotted the median modularity rescaled with the community sizes (Fig. 7). The solid lines represent each chromatin group, including the global median modularity as a reference curve (dashed). These curves show that A–C communities (“active chromatin”) have higher modularity than chromatin group D (“inactive chromatin”) and the entire Hi-C network. This implies that A–C communities partition the network better than the D communities. To complement Fig. 7, we show the 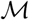 variability in Fig. S10.

**FIG. 7.**
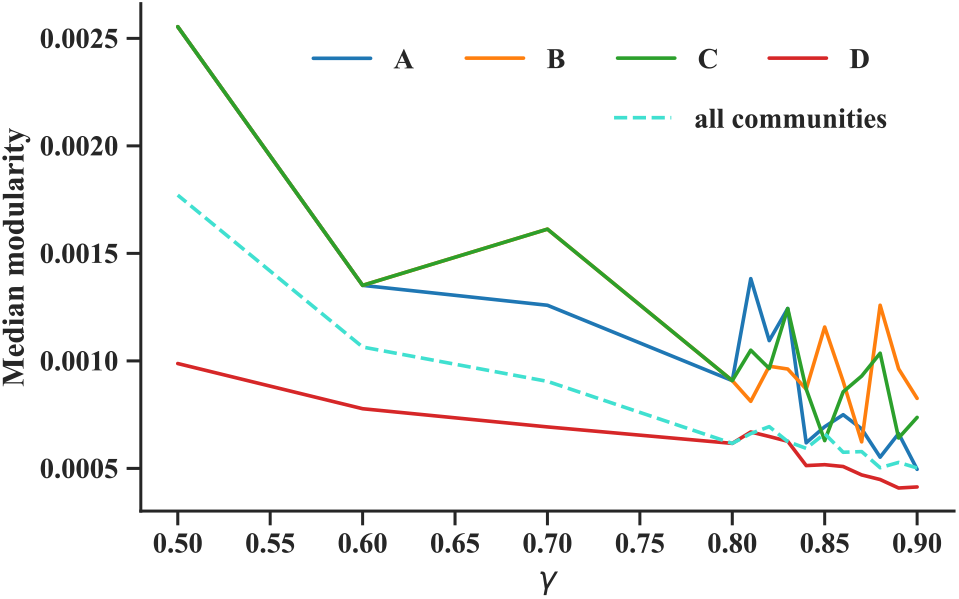
Median community modularity (rescaled with community size) for four chromatin groups A—D (solid lines) across different scale parameters *γ*. The dashed line shows the median for the entire network. A, B, and C communities have higher modularity than D and the whole network.

In addition to forming tighter node clusters, A–C communities tend to nest with each other (as shown in Fig. 4 and exemplified on a subset of domains in Supplementary Material, Fig. S7). These findings argue that active chromatin is hierarchical. At least it is more hierarchical than the D communities that form less convincing communities with substantial random nestedness *N_ij_*. As we concluded from our simple folding model (Sec. III.D), random nesting breaks ideal hierarchies.

## IV. DISCUSSION

In this paper, we have mapped out the semi-hierarchical organization of chromatin in human cells. Viewing the Hi-C data as a DNA contact network, we extracted significant 3D structures using the GenLouvain community detection algorithm that allows us to scan seamlessly through different organization scales. Contrasting common assumptions, the communities form non-hierarchical structures, where some organizational levels show substantial randomness. To better understand this result, we developed a model blending hierarchical folding and random contacts. This model reproduces the degree of nestedness we observe in actual data. We also study the nestedness in terms of chromatin states. We uncover that transcriptionally active states tend to nest more with each other and form more distinct 3D communities relative to the chromosome-wide average and inactive or repressed chromatin.

Our results derive from 100 kb Hi-C data. However, our approach is not restricted to any specific resolution or interaction matrix. It can efficiently analyze various chromatin interaction matrices such as single-cell Hi-C (scHi-C^39^), HiCap^40^, HiChIP^41^, and distance matrices^42^. Nevertheless, modifications to the GenLouvain null model may be necessary for some of these scenarios. For instance, if using Hi-C at higher resolution, e.g., 1 kb, numerous contacts appear inside TADs where the contacts decay as a power-law with an exponent of −0. 75, rather than approximately −1^43^.

Next, we use the GenLouvain method to extract 3D communities. However, generating communities this way is a random process, meaning that the nodes’ community membership will differ between realizations even if the scale parameter *γ* is the same. Interestingly, the community–node correlation varies with *γ*, indicating that some communities are more stable than others. This problem exists in most complex networks where multiscale interactions govern the organization^44^. Therefore, depending on algorithm design, two community detection methods focusing on slightly different connectivity features may disagree on the optimal node assembly. This aspect is mainly unexplored for chromosome organization and worth pursuing in future work.

Furthermore, different community detection algorithms may disagree on the hierarchical levels. For instance, the Leiden algorithm^45^ could yield a more strict community hierarchy than GenLouvain (the Leiden algorithm was developed to “solve” the non-hierarchical features of the original Louvain method). One of the authors of this paper tried the Leiden algorithm to study general community inconsistency problems^?^ While studying the effects of other algorithms is a reasonable research direction, it does not invalidate our approach, which is agnostic to the specific algorithm choice.

There are several TAD-finding methods^46^. These can be broadly categorized into feature-based algorithms, clustering methods, and graph-partitioning tools^47^. In this paper, we use a technique that falls into the graph-partitioning category, which includes perhaps the most popular community-detection algorithm based on modularity maximization. For example, Ref.^48^ finds TADs using Louvain but assumes that the background connectivity is a random network under given node degrees (the Newman-Girvan model). In contrast, we use the fractal-globule null model, which better agrees with the empirical distance decay in contact probability in human Hi-C maps. Although the approach is similar to ours— varying *γ* and extracting communities—there are meaningful quantitative differences. For example, the fractal globule model tends to capture widely spread and delocalized 3D communities, while the Newman-Girvan model typically groups contiguous DNA stretches into local communities, like TADs. In addition, another reference combines maximum modularity and Hi-C-like distance decay and extracts communities for different *γ* values^49^. However, they treat TADs as unbroken DNA stretches, not delocalized as we do here. Using polymer simulations^16^, we demonstrated that this generalization partitions spatially close monomers into meaningful 3D communities.

We interpret our data as active chromatin being more hierarchical than inactive chromatin. From a biological standpoint, this has exciting implications. In active chromatin, there is a menagerie of specific proteins, like transcription factors, that coordinate transcription regulation. These proteins interact with chromatin elements at all distances, bringing some in 3D proximity to regulate transcription. These interactions are not random, so they could contribute to shaping the 3D structure toward a perfect hierarchy. While we lack data to validate this hypothesis, we note that our simple folding model requires fine-tuned interactions to create an ideal hierarchical order and that a slight degree of arbitrary nesting causes noticeable deviations.

While specific proteins regulate transcription in active chromatin, inactive chromatin is often epigenetically repressed. The proteins managing epigenetic repression decorate chromatin with chemical tags over large DNA regions (e.g., methylation of specific histone sites). In this respect, epigenetic repression is not relying on characteristic long-range attractions. It is enough if the right chromatin type is close. If so, this idea foreshadows many random contacts manifesting as a broad nestedness distribution, as in Fig. 4, and a less hierarchical structure than active chromatin.

Several lines of evidence indicate that compartments and TADs emerge from distinct mechanisms, like loop extrusion and phase separation, and are not a hierarchy that stems from identical phenomena operating on different scales. Our paper sheds new light on the hierarchical organization resulting from these phenomena. While large sections nest, others segregate, and there is a significant portion of randomness. We anticipate that cross-scale relationships capturing these features will be essential components in future models aiming to reach a deep understanding of the causal mechanisms of chromosome folding. In the short perspective, our results further open questions worthy of research, including the reliability of 3D communities or what biological factors govern the different organization scales, such as DNA-binding proteins, epigenetic marks, or general chromatin types.

## Acknowledgments

We acknowledge financial support from the National Research Foundation (NRF) of Korea (S.H.L. grant Nos. NRF-2021R1C1C1004132 and NRF-2022R1A4A1030660), the Swedish Research Council (L.L. grant Nos. 2017-03848 and 2021-04080), and the Knut and Alice Wallenberg Foundation (grant No. 2014-0018, co-PI: P.S.)

## Footnotes

1 However, we can get *N_ij_* = −1 in rare cases even if there is some domain overlap. This happens when *d_i_* + *d_j_* > *n* in Eqs. (3)–(7).

