## supplementary figures and text for "Mapping the semi-nested community structure of 3D chromosome contact networks"

Dolores Bernenko*

*Department of Physics, Integrated Science Lab, Umeå University, SE-901 87 Umeå, Sweden.*

Sang Hoon Lee†

*Department of Physics and Research Institute of Natural Science, Gyeongsang National University, Jinju 52828, Korea and*
*Future Convergence Technology Research Institute, Gyeongsang National University, Jinju 52849, Korea*

Per Stenberg‡

*EMG, Umeå University, SE-901 87 Umeå, Sweden*

Ludvig Lizana§

*Integrated Science Lab, Department of Physics, Umeå University, SE-901 87 Umeå, Sweden*

(Dated: March 30, 2023)

### I. SUPPLEMENTARY TEXT AND FIGURES

#### A. Communities and domains for chromosomes 3, 5, and 22

In the main text, we show the domains for chromosome 22 In Fig. S1, we show the domain structure for chromosomes 3, 5, and 22. We use these domains to calculate the nestedness $N_{ij}$ in Fig. 4 (main text).

#### B. Community nestedness for chromosomes 3, 5, 10, and 22

In Fig. 4 (main text), we plot community nestedness $N_{ij}$ over four chromosomes (3, 5, 10, and 22). Figure 4 also shows significant versus random community overlaps. Here in Fig. S3, we show the data for each chromosome separately, panels (a)—(d). In addition, we outline below a specific example where we apply Eqs. (3)—(10) (main text) to calculate $N_{ij}$ and the associated $p$-value in chromosome 22.

To calculate $N_{ij}$, we start by organizing domains and communities in a two-layer bipartite graph [Fig. S2(b)]. One layer contains domains, and the other has all communities associated with different $\gamma$ values. Next, we connect these layers with links representing domain-community memberships. We illustrate how we map out these relationships in Fig.S2(a), where the domains form the ring, ordered according to DNA sequence, and the inner black circles indicate two communities. The links illustrate domain-community memberships.

To build the nestedness histograms, we calculate $N_{ij}$ for every community pair, excluding pairs belonging to the same $\gamma$. As an example, we will calculate $N_{ij}$ explicitly for communities $7_{0.9}$ and $4_{0.7}$ using the method we outline in Sec. II.B (main text).

First, we extract four numbers from the bipartite graph [Fig.S2(b)]:

1. Number of domains: $n = 93$,
2. Number of shared domains: $k = 1$
3. Number of domains per community (community degree): $d_i = 7(4_{0.7})$, $d_j = 6(7_{0.9})$,

Notably, chromosome 22 has 96 domains. But we exclude three of them from the analysis ('1', '3', '96'—shown as large white circles) because they do not form any community regardless of $\gamma$. One of these domains is the centromere (domain '1').

Based on the numbers $k$, $n$, $d_i$ and $d_j$, Eq. (3) predicts that the community pair $7_{0.9}$ and $4_{0.7}$ share 0.45 domains under the random null hypothesis. However, the actual number is higher, $k = 1$. To reach full nestedness, these communities should have shared six domains, which is the maximum possible overlap ($\min(d_i, d_j)$). To reach this limit, the communities need $6 - 0.45 = 5.55$ more domains than the random expectation. But it is only $1-0.45 = 0.55$. These numbers give the nestedness $N_{ij} = 0.55/5.55 = 0.099$.

Next, we calculate the $p$-value associated with the nestedness. Using Eq. (8) and (10), we estimate that the probability of sharing at least one domain is 0.38. This value is much higher than the threshold of significance we set to 0.025. Therefore, we conclude that this overlap is insignificant.

*Electronic address:
†Electronic address:
‡Electronic address:
§Electronic address:

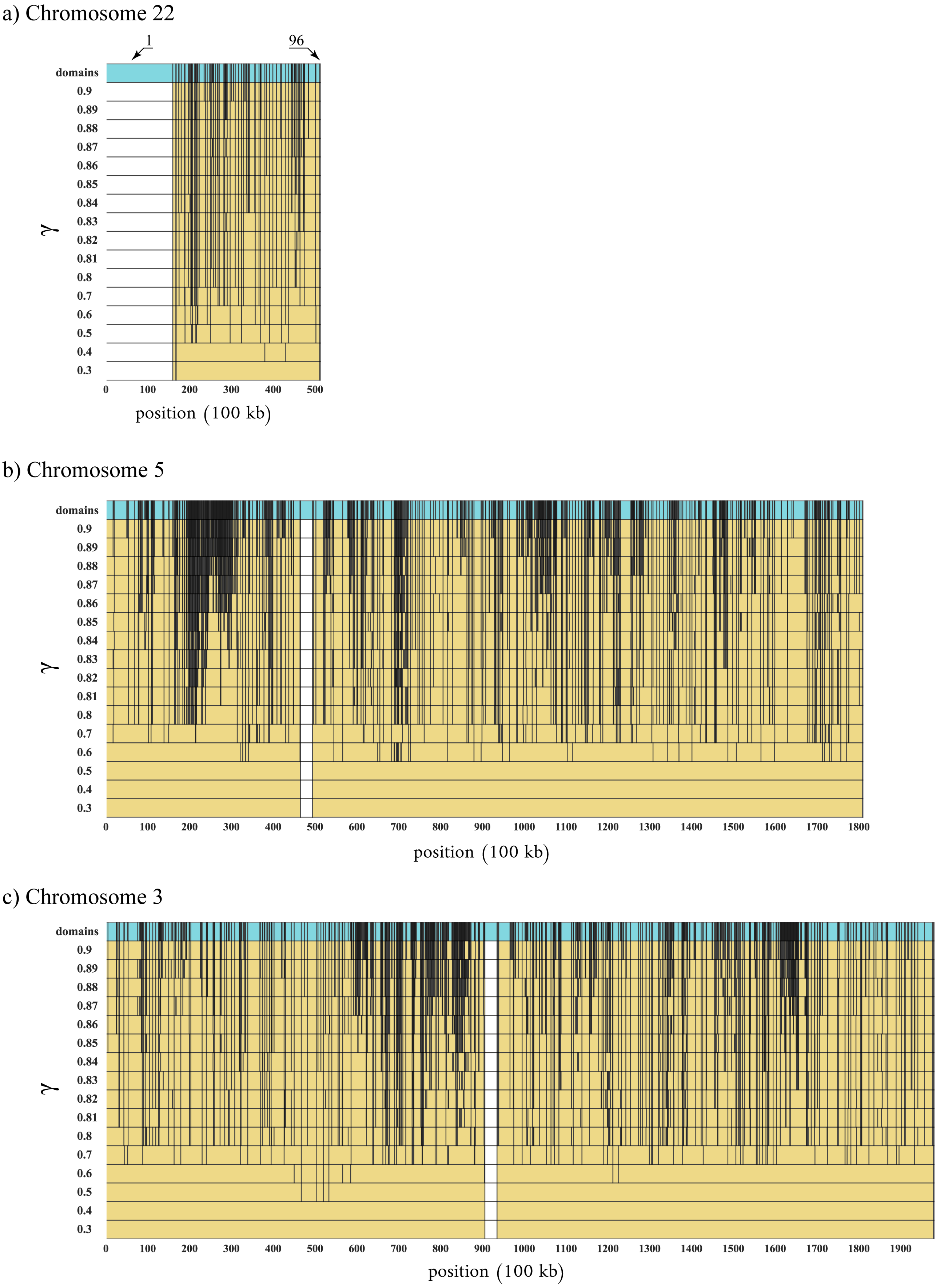


FIG. S1 Division of the Hi-C network into 3D communities and domains across 16 $\gamma$ values for human chromosomess 22(a), 5(b), and 3(c). Each stripe represents a community partition for a single $\gamma$ value. Within each stripe, vertical lines separate two adjacent DNA segments that belong to different communities. The white area shows the centromere. The top turquoise stripes in (a)—(c) show the domains. These are DNA segments that did not break across the shown $\gamma$ range. In (a), we label two domains '1' and '96,' representing the first a last domain along chromosome 22. The domain IDs follow sequential order along the chromosome so that domain 2 is a linear neighbor of domains 1 and 3.

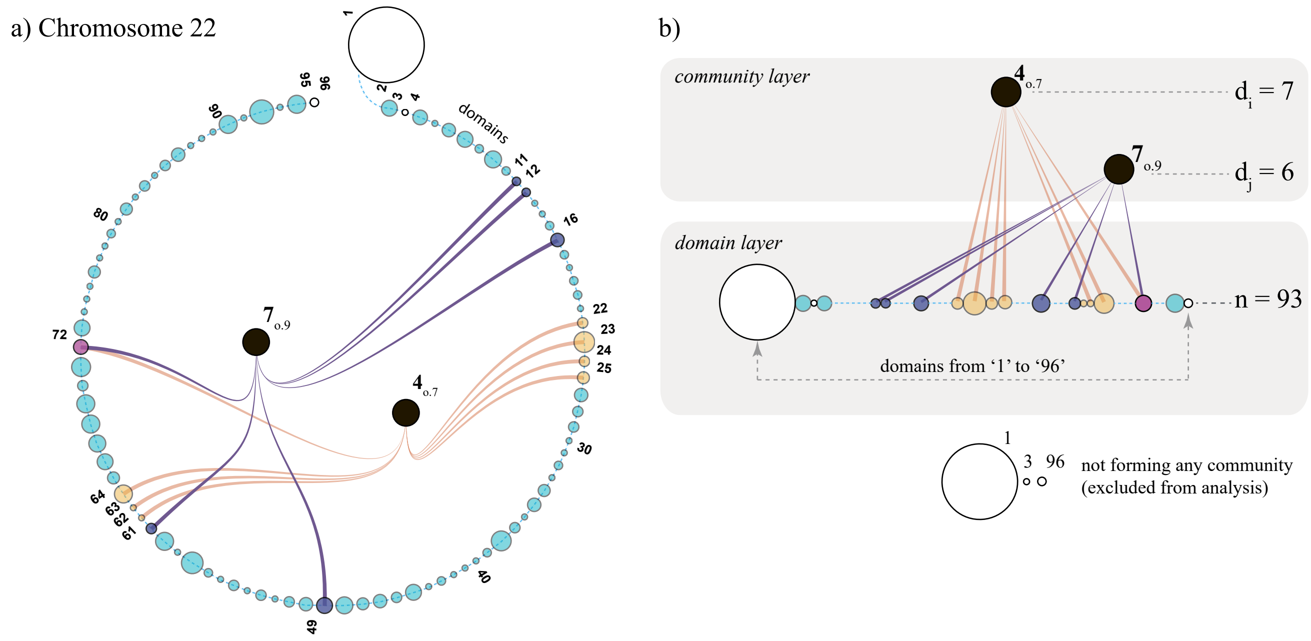


FIG. S2 Community-domain network. (a) We illustrate the domains as a chain of circles starting from '1' and ending with '96.' With colored links, we show domain memberships in two communities: $7_{0.9}$ and $4_{0.7}$. (b) Example of community pair overlap in a bipartite graph. The domains in the bottom layer connect to communities in the upper layer. White circles show domains that do not belong to any community (excluded from the nestedness analysis).

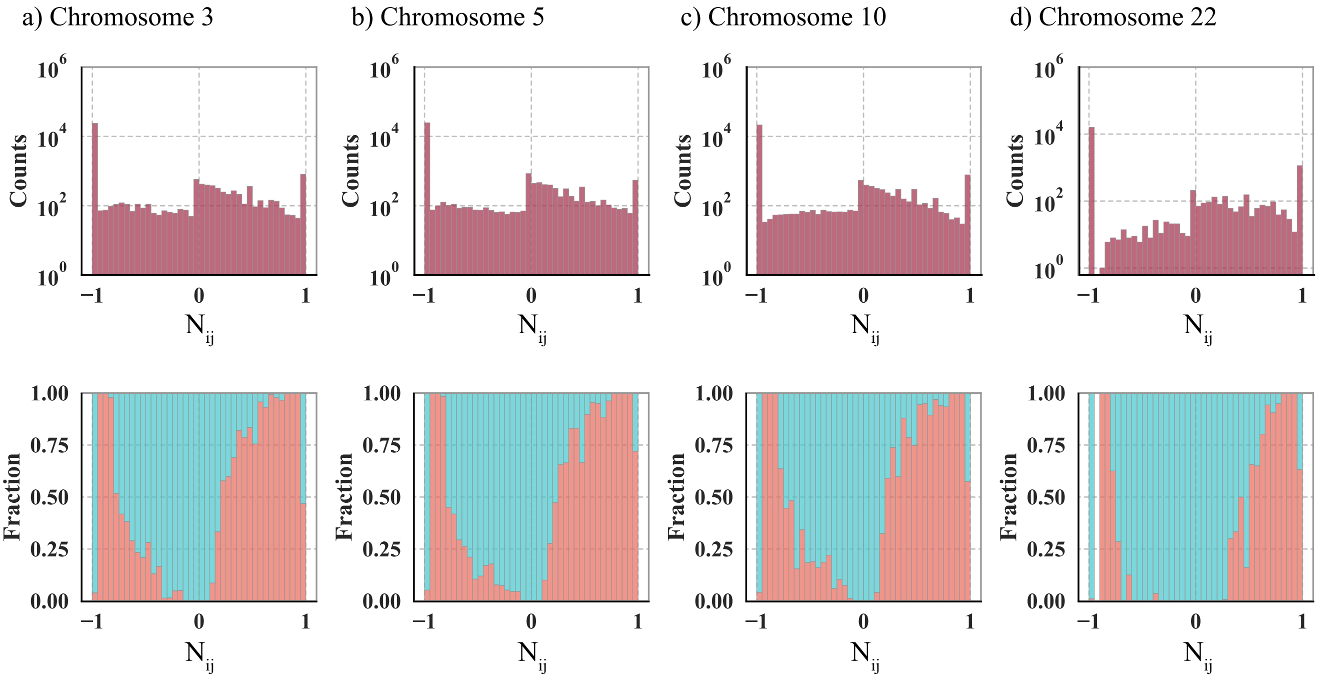


FIG. S3 (top) Nestedness histograms for chromosomes 3, 5, 10, and 22. (bottom) The fraction of $N_{ij}$ scores that are significant (orange) or random (blue-green)

#### C. Characterization of irreducible domains

Figure 2 (main text) shows the division of the chromosome into irreducible domains (top stripe). This figure highlights that domains have different sizes. To understand these sizes better, we construct a letter-value plot [Fig.S4a)], including all irreducible domains depicted along the turquoise stripe in Fig. 2 (main text). While the largest domain size is 30 Hi-C bins, the median is one Hi-C bin (100 kb); 100 kb is identical to the resolution limit of the Hi-C data. If raising $\gamma$ beyond 0.9, the size distribution would have an even higher fraction of one-bin-sized domains. Therefore, to avoid over-partitioning, we select $\gamma = 0.9$ as the upper limit. This allows us to study the chromosome's 3D architecture at a granularity slightly above typical TAD sizes.

#### D. Characterization of structural scales

In Fig. S4b), we depict how the number of communities grows with the scale parameter $\gamma$, where the black line (and symbols) indicate the actual data (chromosome 10). We note that the number of communities grows exponentially (red line) with increasing $\gamma$. We also observe high volatility around this average growth rate for $\gamma > 0.8$. This observation motivates us to sample $\gamma$ more frequently in the range $0.8 \leq \gamma \leq 0.9$ as we expect significant structural rearrangements. To highlight our $\gamma$ choices in Fig. 3 (main text), we mark some data points blue, leaving the others black.

#### E. Nestedness distribution for specific $\gamma$-pairs

In Fig. S5, we show the nestedness distribution ($N_{ij}$) between associated with different $\gamma$ pairs, $\gamma_1$ and $\gamma_2$. In each panel, we fix $\gamma_1$ (0.6, 0.7, 0.8, or 0.9) while varying $\gamma_2$. We illustrate the pair-wise nestedness as stacked $N_{ij}$ histograms. We observe that when $\gamma_1$ is high, the counts are higher than random community overlap ($N_{ij} \sim 0$) since more community pairs are partially nested than segregated [(panels c)(§ and d)].

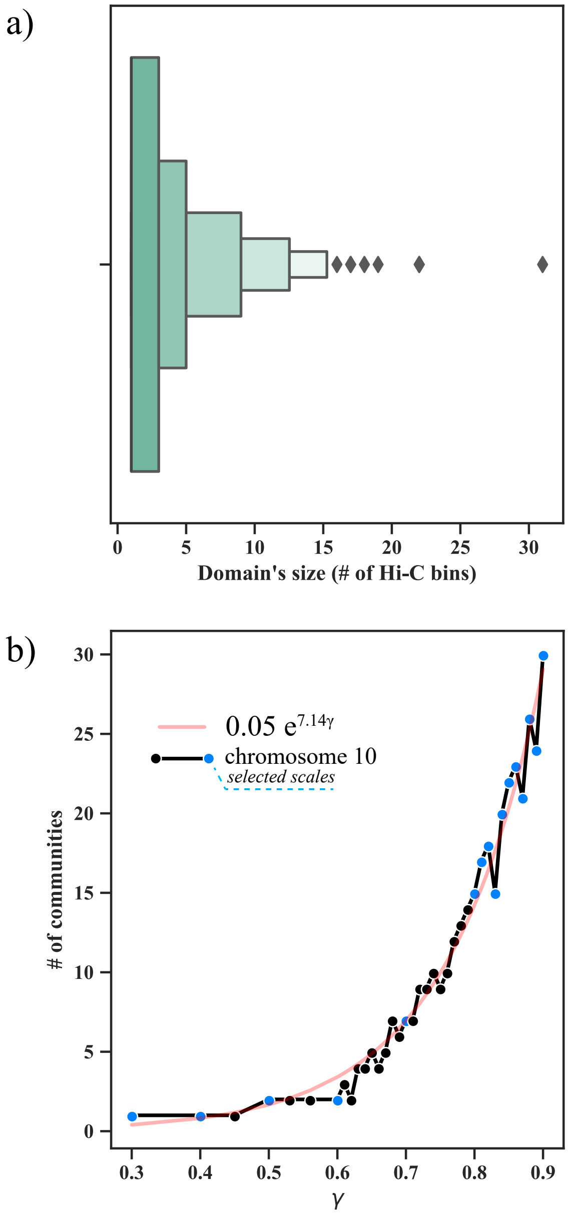


FIG. S4 Characterization of domains and communities (human chromosome 10). (a) Letter-value plot showing the size distribution of irreducible domains. The domain sizes vary between $\sim 1 - 30$ Hi-C bins. The median domain size is 1 Hi-C bin (100 kb). (b) The scale-dependent number of communities (defined by $\gamma$). The number of communities grows exponentially with $\gamma$ (red).

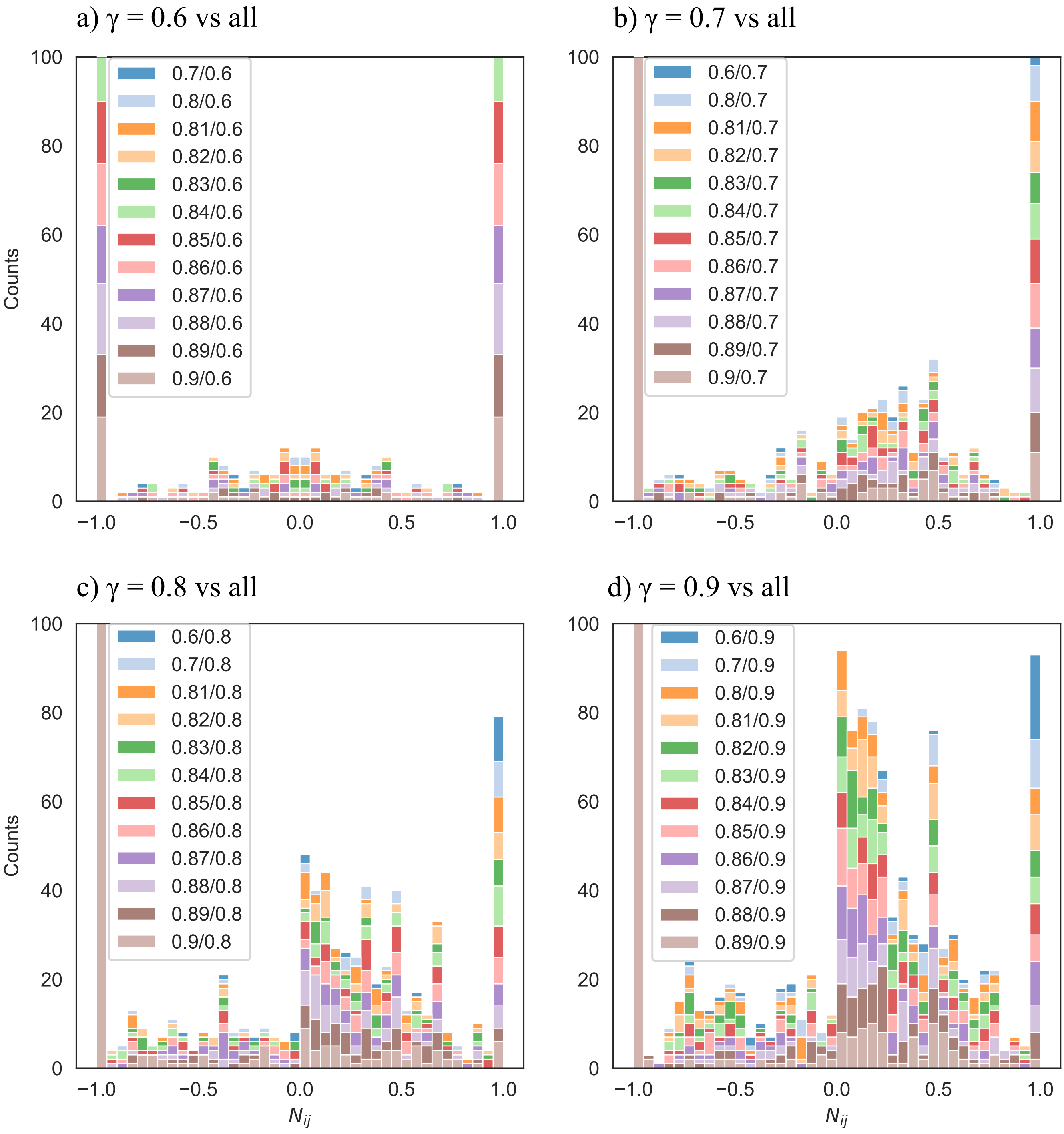


FIG. S5 $N_{ij}$ histograms for individual $\gamma$-pairs (stacked bars with colors specified in legend). To better illustrate range $-1 < N_{ij} < 1$, we truncate the bars at $N_{ij} \pm 1$ if their counts exceed 100. **a)** pairs between $\gamma = 0.6$ and all other $\gamma$. Cross-scale interaction between communities found at $\gamma = 0.6$ and all other communities are fully nested, fully segregated, but also their nestedness is close to random ($-0.5 < N_{ij} < 0.5$); **b)** for $\gamma = 0.7$ and all other $\gamma$. The distribution changes for $N_{ij} \sim 0$), showing that communities tend to be more nested (more counts in the range: $0 < N_{ij} < 0.5$); **c** and **d)** for $\gamma = 0.8$ and 0.9 versus all other $\gamma$. We observe that distribution peaks near $N_{ij} = 0$ when comparing to other distributions.

#### F. Kolmogorov-Smirnov test for nestedness and chromatin types

In the main text, we calculated community nestedness and showed the $N_{ij}$ histograms for all chromatin type combinations (Fig. 6). We found that communities enriched with active chromatin (types A, B, and C) are more nested than those enriched in heterochromatin (type D). In this section, we quantify this observation further by using a standard Kolmogorov-Smirnov test (KS-test) that compares pairwise $N_{ij}$ cumulative distribution functions (CDFs) and calculates a “distance” ($D$). Specifically, we calculate such distances between the AA–DD, BB–DD, and CC–DD distributions.

Figure S6 presents the KS-test analysis. In panel (a), we plot the histograms for distributions AA, BB, CC, and DD (duplicated from Fig. 6, main diagonal). In panel (b), we show the CDFs associated with these distributions and the KS-distance (D) between DD and all other CDFs. Analyzing panel (b), we observed that all CDFs start out at high values (Prob.$>0.6$) due to the peak at $N_{ij}=-1$. This peak indicates complete segregation, which is a common feature for all four histograms. However, the DD distribution is the most segregated: the pink line exceeds all other CDFs for $N_{ij}<0$. Moreover, we note that the AA distribution (green) is the one deviating most from DD. To illustrate their maximum distance, we inserted a vertical bar in panel b) for $N_{ij}<0$ showing $D_{AA}=0.137$ (p-value: $8.1\times10^{-15}$, two-sided).

Next, we analyze the positive $N_{ij}$ range associated with nestedness. We observe that the AA–CC chromatin states are more nested than the DD group because the CDF curves grow faster for $N_{ij}>0.6$ and eventually cross the pink line when $N_{ij}>0.7$. When using the KS test to compare the AA, BB, and CC distributions to DD, we found that only BB and CC were significantly greater than DD. Specifically, the only values with p-value $<0.05$ for the one-sided test were $D_{BB}=0.038$ and $D_{CC}=0.036$.

Overall, we conclude that the KS test supports the notion that chromatin type is strongly associated with the nestedness of communities and that the active genome is more nested than the inactive genome. KS test also shows that the observations are statistically significant.

#### G. Folding pathways and chromatin types

To better understand the biological relevance of folding pathways in Fig. 3b) in the main text, we added colors to highlight the dominant chromatin type of each irreducible domain (Fig. S7). This figure helps us understand whether communities consist of biologically similar domains across different $\gamma$ values. As before, we used the hypergeometric test to quantify the enrichment for each chromatin group. The colors denote:

Red: Enriched in at least one HMM state from groups A, B, or C.

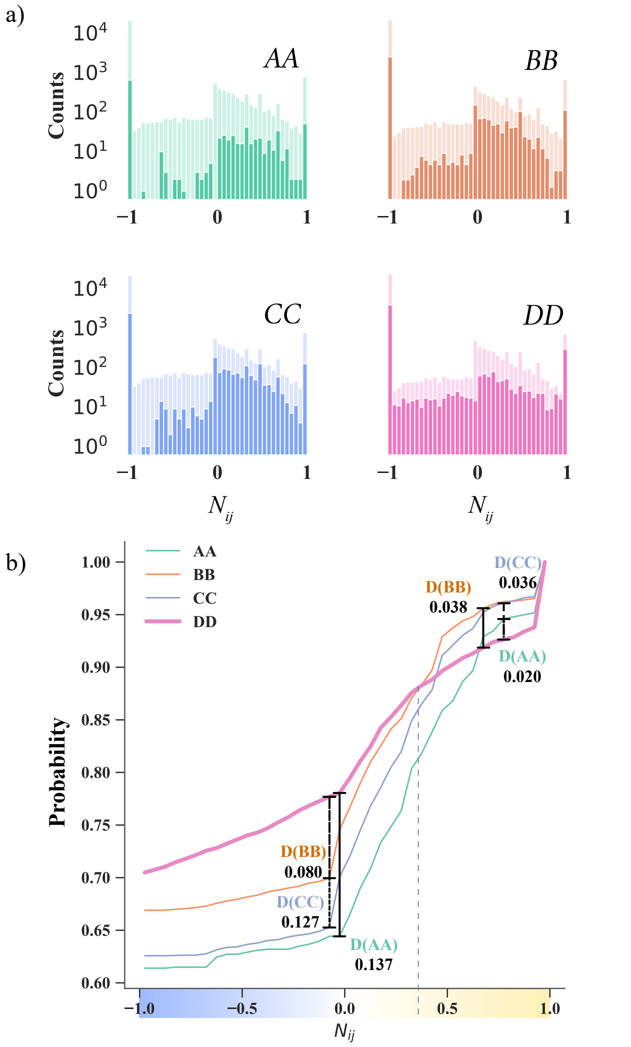


FIG. S6 Nestedness and chromatin type. (a) Nestedness distributions ($N_{ij}$) between communities of the same type (either A, B, C, or D). (b) CDF of $N_{ij}$ distributions. The vertical bars indicate the Kolmogorov-Smirnov distance between 'DD' distribution (thick pink line) and all others.

Blue: Enriched in HMM states from group D.

Green: Enriched in A, B, or C chromatin groups and at least one HMM state from group D.

Gray: No significant enrichment of any chromatin states (p-value $=0.025$).

The figure illustrates how one node (irreducible domain) may be isolated from other nodes having the same chromatin type at some $\gamma$ value but joins them at two other $\gamma$ values, one lower and one higher. For example, we may see this scenario if tracking some of the red circles inside the 3D community labeled 10 ($\gamma=0.9$), that passes through communities 3 and 19 ($\gamma=0.86$), and 5 ($\gamma=0.3$). These irreducible domains assemble in a complex merging-and-splitting behavior outlined

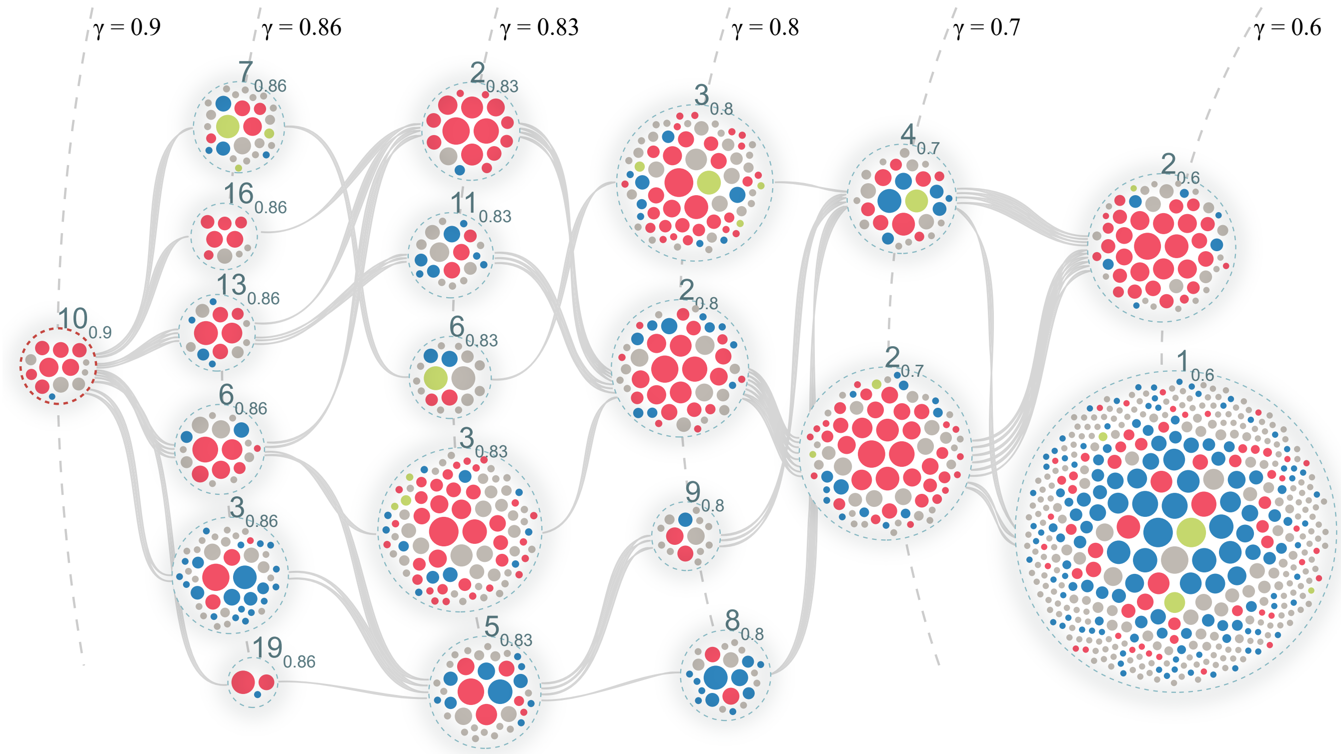


FIG. S7 Rearrangements of chromatin domains across structural scales. Color codes represent their dominant chromatin types. Red domains are enriched in A/B/C states, blue domains in the D-group, green domains in a combination of A/B/C and D-group states, and gray domains show no significant enrichment (p-value set at 0.025). The domain content of communities varies, with some dominated by active chromatin types and others suppressed, similar to the well-known A/B compartmentalization. As resolution increases from $\gamma = 0.6$ to $\gamma = 0.9$, domains reorganize into a community dominated by A/B/C chromatin. At intermediate scales, domains preferentially exchange between structural scales while maintaining biological similarity.

above. Overall, the 3D communities tend to harbor irreducible domains having the same color, suggesting that they typically enrich domains with identical chromatin types. While we do not have enough data to suggest specific molecular mechanisms for this behavior, our data support a non-nested folding scheme.

### H. Optimal Q-parameter

Searching for the optimal Q parameter for the model described in Section D: "Modeling non-nested chromosome folding", we scan Q in the range from 0.01 to 0.5, aiming to minimize the distance between $N_{ij}$ distribution from the real data and the model. As the distance metric, we choose the Kolmogorov-Smirnov's (KS) distance that reports the maximal difference between two cumulative distribution functions (CDF) in a two-sided statistical test. Before measuring the KS distance, we omit the extreme $N_{ij}$ values ($N_{ij} \pm 1$) from all distributions as they are the strongest features that bias the search for the optimal Q.

In Fig. S8(a), we plot CDF distributions for the data (chromosome 10, black thick line) and various Q parameters. All CDF plots have no counts at $N_{ij} \pm 1$. The CDF for the optimal Q-parameter is shown as a thick blue line.

Fig. S8(b) shows the largest distance between the CDF of the real data and Q. We observe that $Q_{opt} = 0.3$. Reshuffling domains with a probability of 30 percent at each structural scale, we obtained similar to the real data distribution around $N_{ij} = 0$, representing the random overlap. We visualize the CDF of our model at $Q = 0.3$ and the real data in Fig. S8(c, d).

### I. Community modularity and chromatin type

In Fig. S9, we show that community modularity depends on size. We use this size-dependence to rescale the modularity to facilitate comparisons. We also show that the rescaled modularity varies among chromatin types (Fig. S10).

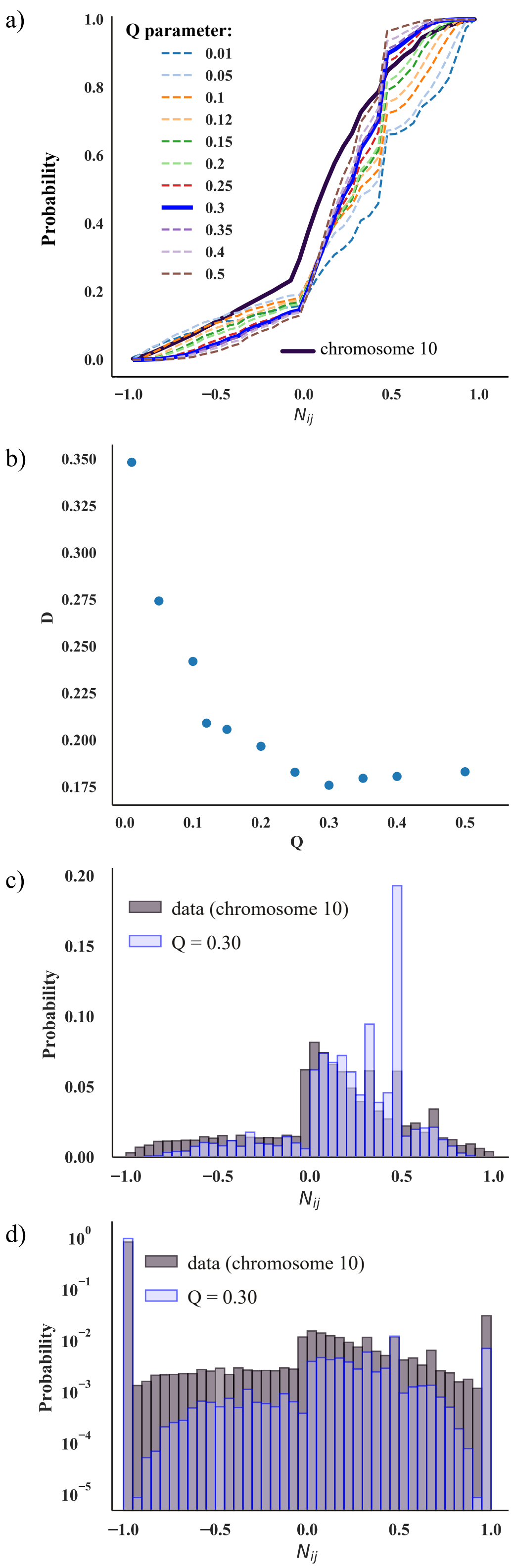


FIG. S8 Optimal $Q$ analysis for chromosome 10. (a) CDF for nestedness score for the real data (black thick line) and models associated with varying $Q$. In all CDFs, the extreme $N_{ij} = \pm 1$ values are excluded. (b) Kolmogorov-Smirnov distance, $D$, versus the reshuffling parameter $Q$. We find the minimal distance when $Q = 0.3$. This represents the optimal $Q_{rmopt.}$. (c) Normalized $N_{ij}$ distributions for chromosome 10 and $Q = 0.3$ with $N_{ij} = \pm 1$ removed from the plot. (d) Same as panel c), but including all $N_{ij}$ counts. We use log-scaled axes to fit the $\pm 1$ peaks.

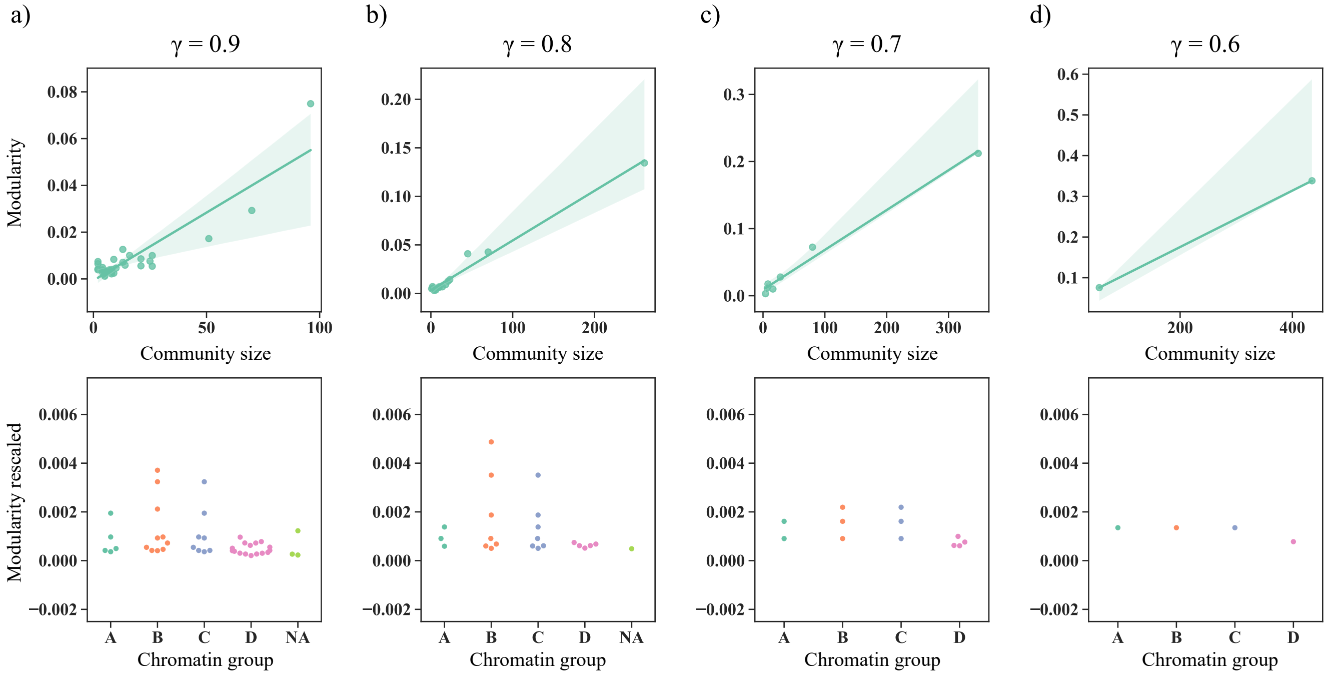


FIG. S9 Community modularity for four $\gamma$ values: (a) $\gamma = 0.9$, (b) $\gamma = 0.8$, (c) $\gamma = 0.7$, (d) $\gamma = 0.6$. The top panels show a linear regression fit between community size (number of domains) and modularity (Eq. (1), main text); We observe a nearly linear relationship. The bottom panels show the community modularity rescaled with the community size. We define the A—D chromatin groups in Sec. II (main text). The 'NA' group represents communities that are not enriched in any chromatin group.

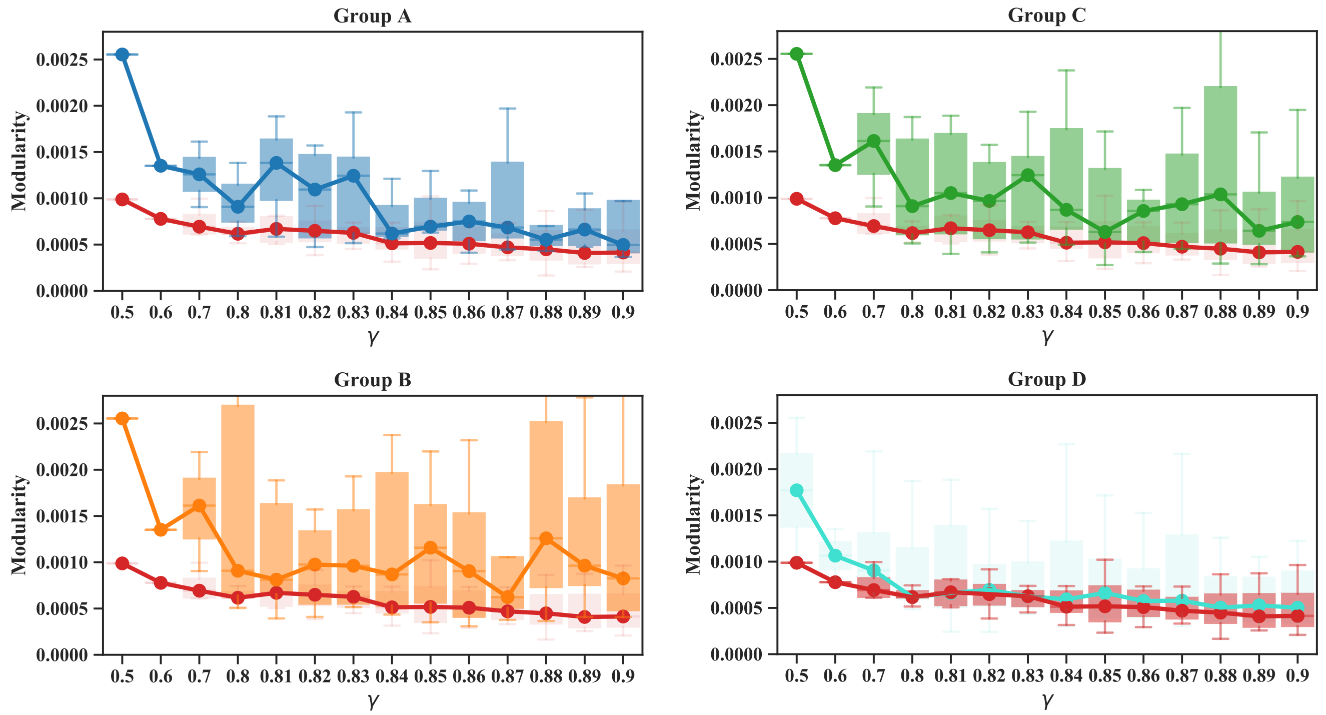


FIG. S10 Community modularity (rescaled with community size) for chromatin groups A—D across different scale parameters $\gamma$. The top four plots visualize community modularity as bar plots, where values of median modularity are connected across $\gamma$. In bar plots, we compare groups A (blue), B (yellow), and C (green) with group D (red bar plot). However, the D group we compare with the modularity of all communities (light blue plot).
